# Delaying or Delivering: Identification of novel NAM-1 alleles which delay senescence to extend grain fill duration of wheat

**DOI:** 10.1101/2020.09.23.307785

**Authors:** Elizabeth A. Chapman, Simon Orford, Jacob Lage, Simon Griffiths

## Abstract

Senescence is a complex quantitative trait under genetic and environmental control, involving the remobilisation of resources from vegetative tissue into grain. Delayed senescence, or ‘staygreen’ traits, are associated with conferring stress tolerance, with extended photosynthetic activity hypothesised to sustain grain filling. The genetics of senescence regulation are largely unknown, with senescence variation often correlated with phenological traits. Here, we confirm staygreen phenotypes of two *Triticum aestivum* cv. Paragon EMS mutants previously identified during a forward genetic screen and selected for their agronomic performance, similar phenology and differential senescence phenotypes. Through grain filling experiments, we confirm a positive relationship between onset of senescence and grain fill duration, reporting an associated ∼14 % increase in final dry grain weight for one mutant, *P* < 0.05. Recombinant Inbred Line (RIL) populations segregating for senescence were developed for trait mapping purposes, and phenotyped over multiple years under field conditions. Staygreen traits were mapped using exome-capture enabled bulk segregant analysis (BSA), whereupon senescence was quantified, and metrics compared to qualify senescence traits and aid RIL selection. Using BSA we mapped our two staygreen traits to two independent, dominant, loci of 4.8 and 16.7 Mb in size encompassing 56 and 142 genes. Combining single marker association analysis with variant effect prediction, we identified SNPs encoding self-validating mutations located in *NAM-1* homoeologues and propose these as gene candidates.

**Summary:** Applying an exome-capture enabled Bulk Segregant Analysis, we identified novel *NAM-1* alleles underpinning two staygreen traits associated with enhancing grain fill duration and grain weight

## Introduction

Monocarpic senescence is the terminal stage in wheat development, wherein 80% of leaf nitrogen is remobilised into developing grain (Buchanan-Wollaston, 2007). Genetic regulation of senescence involves significant transcriptional reprogramming enabling timely reallocation of resources. Plants with delayed senescence are described as ‘staygreen’, with cosmetic types resulting from impaired chlorophyll catabolism (Thomas and Ougham, 2014). Functional staygreen phenotypes are associated with enhanced or extended photosynthetic activity, conferring tolerance to heat, drought and low nitrogen stress in multiple crops (Gregersen *et al.*, 2013; Thomas and Ougham, 2014).

Correlations between green canopy and grain fill duration of r = 0.16 to 0.7, *P* < 0.01 (Pinto *et al.*, 2016; De Souza Luche *et al.*, 2017) support the potential breeding utility of staygreen traits. During grain fill Gelang *et al*. (2000) estimates the rate of grain weight increase is 0.96 to 1.25 mg day^-1^, with studies by Adu *et al*.(2011), Kitonyo *et al*. (2017) and Voss-Fels *et al.*, (2019) suggesting staygreen traits have been selected over the last 50 years to sustain grain number improvement. For a *Triticum durum* cv. Trinkaria EMS mutant a 10-day delay in onset of senescence contributed to a 10-12 % and 20 % increase in thousand grain weight (TGW) and yield, respectively (Spano *et al.*, 2003). Under terminal stress, Kumari *et al*. (2007) confirmed the association between grain fill extension, TGW and yield improvement, with staygreen phenotype conferring a yield advantage. Modelling of wheat ideotypes using 2050 climate predictions weights staygreen traits highly, estimating associated yield benefits of 10-37 % for certain Europe regions (Senapati *et al.*, 2019).

Phenology can confound senescence trait dissection (Bogard *et al.*, 2011; Camargo *et al.*, 2016). For example, of the stable senescence QTLs identified by Bogard *et al*. (2011), Naruoka *et al*. (2012) and Xie *et al*. (2016) several controlled anthesis date, encompassing genes *Ppd-D1, Vrn-B1* and *Vrn-D3*. However, Verma *et al*. (2004), Naruoka *et al*. (2012) and Xie *et al*. (2016) reported phenology-independent senescence QTLs located on chromosomes 2DL, 2B and 4A, with Naruoka *et al*. (2012) validating their QTL for ‘Green leaf area duration after heading’ using near isogenic lines. At the gene level, *NAM-B1* and *NAC-S* are known wheat senescence regulators (Uauy *et al.*, 2006a; Zhao *et al.*, 2015). Both genes belong to the plant specific NAC transcription factor family, members of which are involved in hormone signalling, developmental pathways and stress response (Olsen *et al.*, 2005; Podzimska-Sroka *et al.*, 2015).

Induced mutations are a vital source of novel variation, with 2250 mutant derived crop varieties released since 1940 (Ahloowalia *et al.*, 2004). In wheat, exome capture has been used to detect such variation, reducing genetic complexity associated with its large genome through sequencing the gene-encoding 2 % (Henry *et al.*, 2014; Krasileva *et al.*, 2017). Genetic characterisation of TILLING populations and diverse material (Krasileva *et al.*, 2017; Alaux *et al.*, 2018) increases their applicability for use in forward and reverse genetic screens or functional characterisation, accelerating gene discovery.

Here, we use exome capture enabled BSA to map novel staygreen alleles underpinning delayed senescence phenotypes of two independent *Triticum aestivum* cv. Paragon EMS mutants. We confirm results of the initial forward screen, and the relationship between onset of senescence and grain fill duration. Following repeated phenotypic assessment of segregating RILs, we reduced senescence from a quantitative to qualitative trait enabling construction of staygreen and non-staygreen bulks. We used mapping-by-sequencing to identify two independent loci located on chromosomes 6A and 6D. Using variant effect prediction and single marker association analysis we refined these identified regions to likely gene candidates. Here, results converged upon self-validating, dominant, mutations in homoeologous copies of known senescence regulator *NAM-B1* (Uauy *et al.*, 2006a).

## Results

### Identification of Staygreen Mutants

6500 M_5:6_ generation *Triticum aestivum* cv. Paragon EMS mutant lines were phenotyped under field conditions at JIC between 2006 and 2007 (data not shown). Approximately 1.2 % (∼80%) of lines displayed differential senescence phenotypes, with staygreen phenotypes of 1189a and 2316b unconfounded by heading-date variation (Figure 1A). To confirm these results detailed phenotyping of 1189a and 2316b was conducted between 2016 and 2018, identifying staygreen phenotypes are characterised by environmentally stable delays in onset of senescence. Time-course leaf and peduncle senescence profiles of 1189a and 2316b differed significantly compared to cv. Paragon, *P* < 0.05 (Figure 1B, Supplemental Figure S1). Compared to cv. Paragon leaf senescence of 1189a and 2316b is initiated 6-10 and 3-4 days later, respectively, whilst senescence rate is unaffected (Table 1), with pattern of peduncle senescence similar (Figure 1B, Supplemental Figure S1).

**Table 1.** Senescence Pairwise Comparison (P-value, Mutant vs. cv. Paragon) Results of Tukey-post hoc tests comparing overall leaf and peduncle time-course senescence. Date of onset of leaf senescence and days difference relative to cv. Paragon in parentheses. Variation in heading date totalled 0 to 1 days. Peduncle senescence went unscored in 2016

|  | 1189a |  |  | 2316b |  |  |
| --- | --- | --- | --- | --- | --- | --- |
| Year | Onset (dd/mm) | Leaf | Peduncle | Onset (dd/mm) | Leaf | Peduncle |
| 2016 | 29/07 (+ 10) | 0.11 | - | 23/07 (+ 4) | 0.0005 | - |
| 2017 | 06/07 (+ 6) | < 0.0001 | < 0.0001 | 04/07 (+ 4) | < 0.0001 | 0.002 |
| 2018 | 07/07 (+ 6) | < 0.0001 | < 0.0001 | 03/07 (+ 3) | 0.08 | 0.048 |

**Figure 1.**
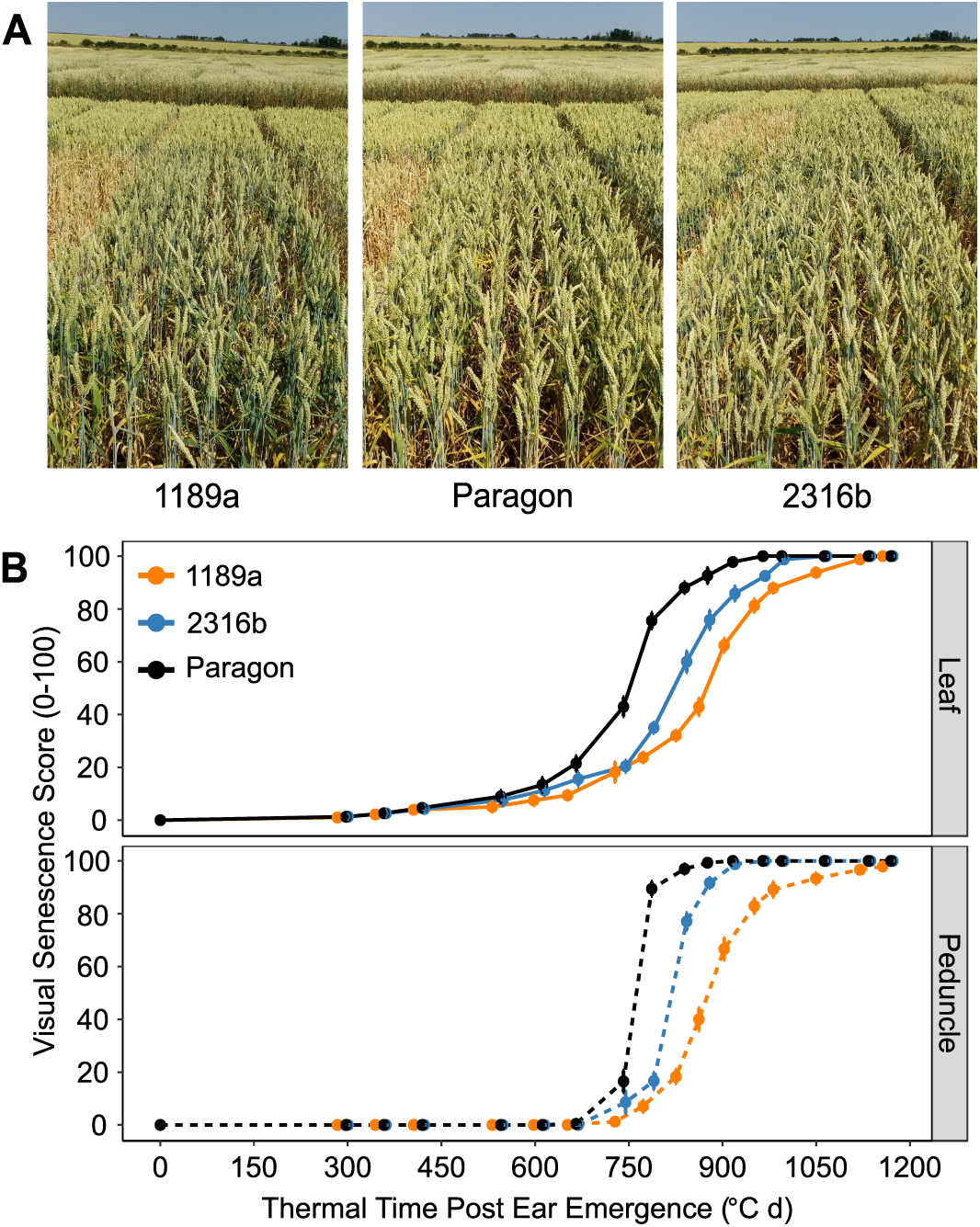
Identification and confirmation of two staygreen mutants. **(A)** Visual differences in senescence of 1189a (right) and 2316b (left) compared to cv. Paragon (centre) relate to variation in onset. Plots photographed 05/07/2018, GS55 within ± 1 day. **(B)** Progression of flag leaf (top) and peduncle (bottom) senescence of 1189a, 2316b and cv. Paragon in 2017. Senescence was scored visually using a 0-100 scale 2-4 times per week from ear emergence (GS55), with scoring dates converted to thermal time (day °C). Mean ± SEM, n ≥ 12. Lines shown, cv. Paragon (black), 1189a (orange), 2316b (blue).

### Extended Grain Fill Duration of Staygreens

Maintenance of green leaves has been associated with extending grain fill duration and grain weight enhancement (Wiegand and Cuellar, 1981; Gelang *et al.*, 2000; Bogard *et al.*, 2011). To test this, we recorded grain weight and moisture content to determine grain fill duration of 1189a, 2316b and cv. Paragon in 2017 and 2018.

Grain moisture content declined more slowly for 1189a and 2316b compared to cv. Paragon, *P* < 0.0001 (Figure 2A & C, Supplemental Figure S3). 42 days after anthesis (daa) grain moisture content of our staygreens was 17 to 20 % greater compared to cv. Paragon, *P* ≤ 0.007 (Figure 2A & C, Supplemental Figure S3, Supplemental Table S1). Subsequently, grain moisture content of 1189a remained elevated, *P* < 0.001, suggesting an extended time to reach grain maturity (Figure 2A, Supplemental Figure S3). Between 48 to 52 daa differences in grain moisture content between 2316b and cv. Paragon were not significant, with time to grain maturation similar, *P* > 0.2 (Figure 2C, Supplemental Figure S3). Relative to cv. Paragon, differences in grain moisture for 1189a and 2316b were significant at 4 and 3 timepoints, respectively (Supplemental Table S1), reflecting the differences in onset of senescence (Table 1, Supplemental Figure S2, Supplemental Figure S3). Grain fill extensions observed for our staygreens did not consistently increase final dry grain weight. Dry grain weight accumulation for 2316b matched cv. Paragon in both years, *P* > 0.1 (Figure 2D, Supplemental Figure S3), but was greater for 1189a in 2018, *P* < 0.001 (Figure 2B). In 2017, between 37 and 47 daa dry grain weights for 1189a and 2316b were greater relative to cv. Paragon, *P* < 0.05 (Supplemental Figure S3), but these differences did not contribute to a significant increase in final grain weights on 23^rd^ July, *P* > 0.05 (Supplemental Table S1). Conversely, for 1189a the significant differences in dry grain weight recorded between 42 and 48 daa, *P* < 0.01, did translate to an 11 to 14.4 % increase in final dry grain weight, *P* < 0.001 (Figure 2B, Supplemental Table S1).

**Figure 2.**
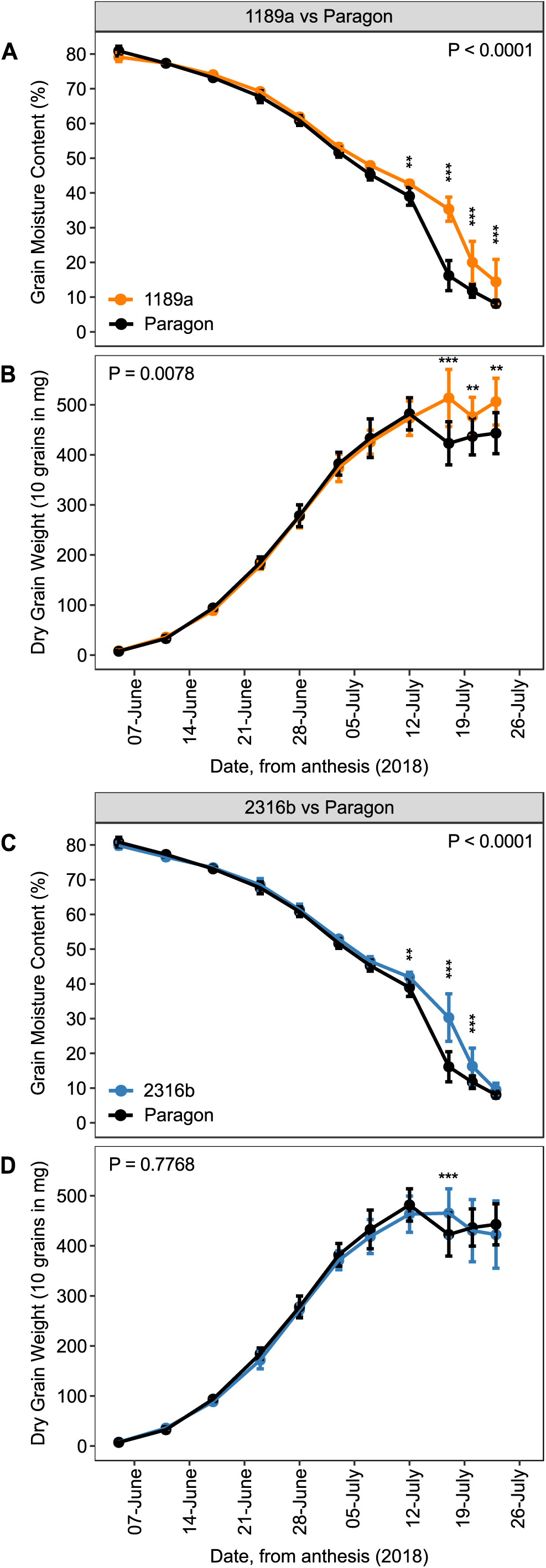
Grain filling dynamics of 1189a and 2316b are altered compared to cv. Paragon. Grain fill duration, determined by the reduction in grain moisture content, is longer for 1189a **(A)** and 2316b **(C)**, *P* < 0.0001. Grain filling extension is associated with greater dry grain weight accumulation and final grain weight recorded for 1189a **(B)**, *P* < 0.001, but not 2316b **(D)**, *P* > 0.2. In 2018, grain moisture content(%) and dry weight(10 grains in mg) were recorded at 3 to 6-day intervals starting from anthesis. Mean ± SD, n = 4-5, 2 plots per line. Printed P-values represent overall differences (ANOVA). Pairwise differences indicated at corresponding time points, *P*: * < 0.05, ** < 0.01, *** < 0.001(Tukey post-hoc test). Corresponding senescence profiles plotted in Supplemental Figure S2. Grain filling experiment was repeated in 2018 following results obtained in 2017(Supplemental Figure S2), with Supplemental Table 3 reporting statistics for both.

### Identifying Senescence Extremes

To genetically map our staygreen traits we adopted a complexity reduction BSA approach. Repeated in-field phenotyping of Paragon x 1189a and Paragon x 2316b RILs enabled their accurate classification into senescence types. For each experiment senescence profiles of individual RILs were quantified by deriving senescence metrics. RILs were considered ‘non-staygreen’ or ‘staygreen’ based on whether their phenotypic scores were lower or higher when compared to cv. Paragon, or the staygreen parent. RILs for which senescence metrics fell between parental values were classified manually based on metric concordance. To assess phenotypic stability of RILs inter-year comparisons were performed, with bulk selections guided be a minimum of one, or two, years of data for 1189a and 2316b, respectively. Details of RIL classification and bulk selections supplied in Supplementary Material 2.

**Table 2.**
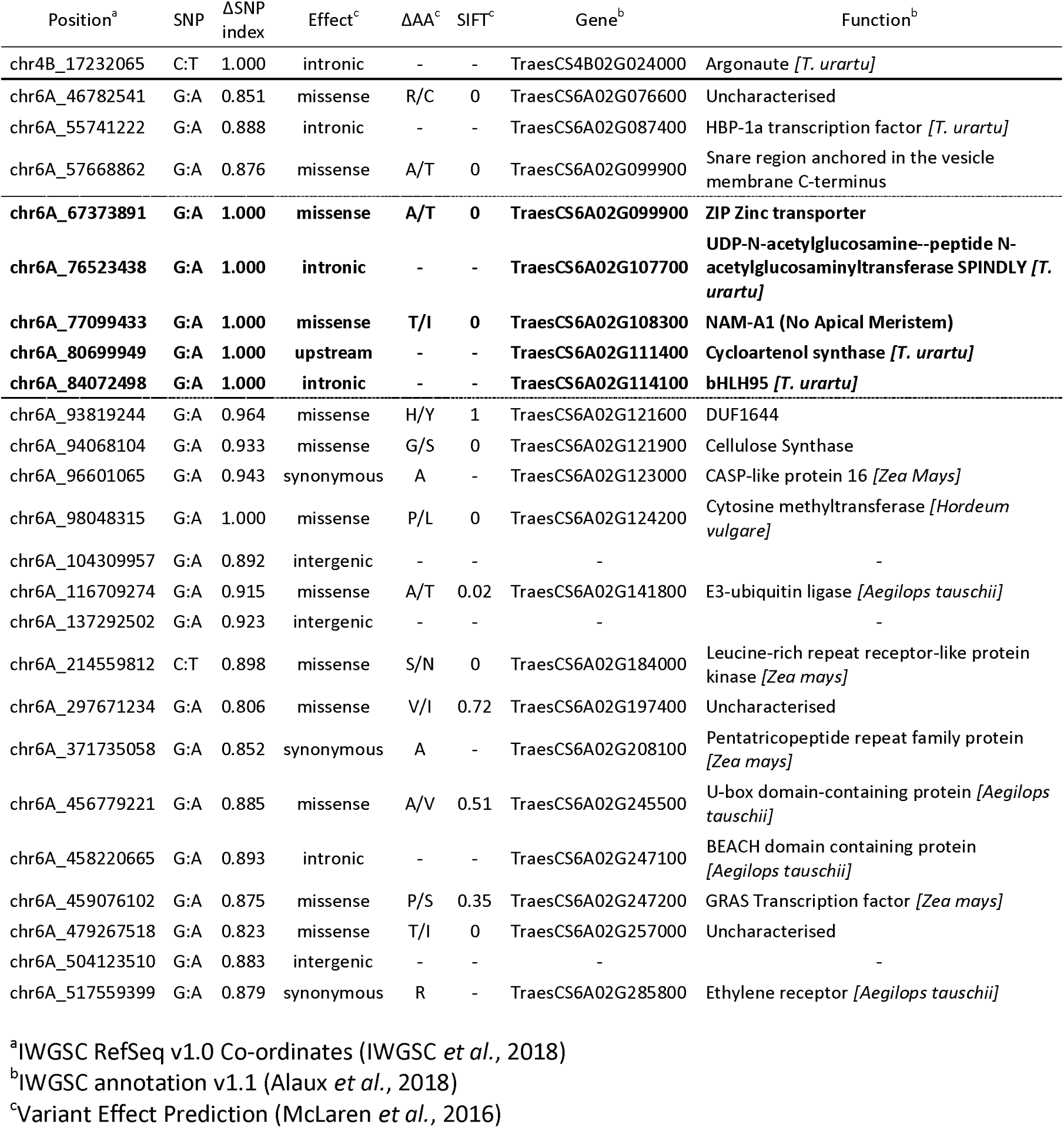
Bulk Segregant Analysis maps the 1189a allele to a chromosomal region on 6A encoding a mutation in NAM-A1.

**Table 3:** Bulk Segregant Analysis maps the 2316b allele to a chromosomal region on 6D encoding a mutation in NAM-D1.

| Position<sup>a</sup> | SNP | ΔSNP index | Effect<sup>c</sup> | ΔAA<sup>c</sup> | SIFT<sup>c</sup> | Gene<sup>b</sup> | Function<sup>b</sup> |
| --- | --- | --- | --- | --- | --- | --- | --- |
| chr2A_732959676 | C:T | 1.000 | intergenic | - | - | - | - |
| chr2D_15472677 | C:T | 1.000 | missense | A/T | 0.38 | TraesCS2D02G042900 | NBS-LRR resistance like protein [*Hordeum vulgare*] |
| chr3A_8326626 | C:T | 0.833 | intronic | - | - | TraesCS3A02G009200 | NB-ARC domain |
| chr3B_12565097 | C:T | 0.875 | intergenic | - | - | - | - |
| chr3D_588999854 | G:A | 0.833 | splice region, synonymous | - | - | TraesCS3D02G498000 | Protein tyrosine kinase |
| chr6D_36628312 | G:A | 0.909 | 3' UTR | - | - | TraesCS6D02G072100 | PR17c precursor [*Hordeum vulgare*] |
| chr6D_42295410 | C:T | 1.000 | intergenic | - | - | - | - |
| chr6D_50958137 | G:A | 0.875 | downstream | - | - | TraesCS6D02G085600 | Uncharacterised |
| **chr6D_57044844** | **C:T** | **1.000** | **intergenic** | - | - | - | - |
| **chr6D_60487321** | **C:T** | **1.000** | **missense** | **G/E** | **0** | **TraesCS6D02G096300** | **NAM-D1 No Apical Meristem** |
| **chr6D_65341694** | **C:T** | **1.000** | **missense** | **G/S** | **0.56** | **TraesCS6D02G101900** | **F-box Domain** |
| chr6D_72022762 | C:T | 0.960 | intronic | - | - | TraesCS6D02G107900 | SNF2 Family N-terminal domain |
| chr6D_79076045 | G:A | 0.909 | missense | A/T | 0.32 | TraesCS6D02G112400 | Diaminopimelate decarboxylase, chloroplastic [*T. urartu*] |
| chr6D_84222670 | G:A | 0.929 | downstream | - | - | TraesCS6D02G119000 | Tyrosine-sulfated glycopeptide receptor 1-like [*Aegilops tauschii*] |
| chr6D_86733052 | G:A | 1.000 | intronic | - | - | TraesCS6D02G122300 | GDP-fucose protein O-fucosyltransferase |
| chr6D_110872402 | G:A | 0.756 | synonymous | - | - | TraesCS6D02G141300 | FGGY carbohydrate kinase domain-containing protein [*Aegilops tauschii*] |
| chr6D_119464714 | G:A | 0.755 | intergenic | - | - | - | - |
| chr6D_139310981 | G:A | 0.817 | intronic | - | - | TraesCS6D02G161000 | Lycopene beta cyclase [*T. urartu*] |
| chr6D_140102996 | G:A | 0.833 | intronic | - | - | TraesCS6D02G161800 | Aminotransferase [*Zea mays*] |
| chr6D_148079666 | G:A | 0.790 | missense | P/S | 0.01 | TraesCS6D02G166500 | Glycosyltransferase [*Aegilops tauschii*] |
| chr6D_264681596 | G:A | 0.815 | missense, splice region | G/D | 0 | TraesCS6D02G192000 | Hypothetical methyltransferase |
| chr6D_292379919 | G:A | 0.941 | intronic | - | - | TraesCS6D02G206900 | S1 RNA Binding Domain |
| chr6D_352357451 | C:T | 1.000 | 5' UTR | - | - | TraesCS6D02G248900 | - |
| chr6D_363004043 | G:A | 1.000 | synonymous | - | - | TraesCS6D02G256800 | GPI-anchored protein [*Aegilops tauschii*] |
| chr7D_8178083 | C:T | 0.833 | synonymous | - | - | TraesCS7D02G018200 | NBS-LRR type resistance |
| chr7D_583986610 | G:A | 1.000 | intronic | - | - | TraesCS7D02G470700 | - |

<sup>a</sup>IWGSC RefSeq v1.0 Co-ordinates (IWGSC *et al.*, 2018)
<sup>b</sup>IWGSC annotation v1.1 (Alaux *et al.*, 2018)
<sup>c</sup>Variant Effect Prediction (McLaren *et al.*, 2016)

Senescence progression was highly dynamic between years, and when classifying and selecting RILs the discriminatory power of senescence metrics varied. In 2017, mean peduncle senescence scores were most discriminative, opposed to duration of leaf senescence in 2018 (Figure 3). The metric TT70 consistently discriminated senescence variation, with large differences always visible between parental lines and segregating RILs (Supplemental Figure S4, Supplemental Figure S5).

**Figure 3.**
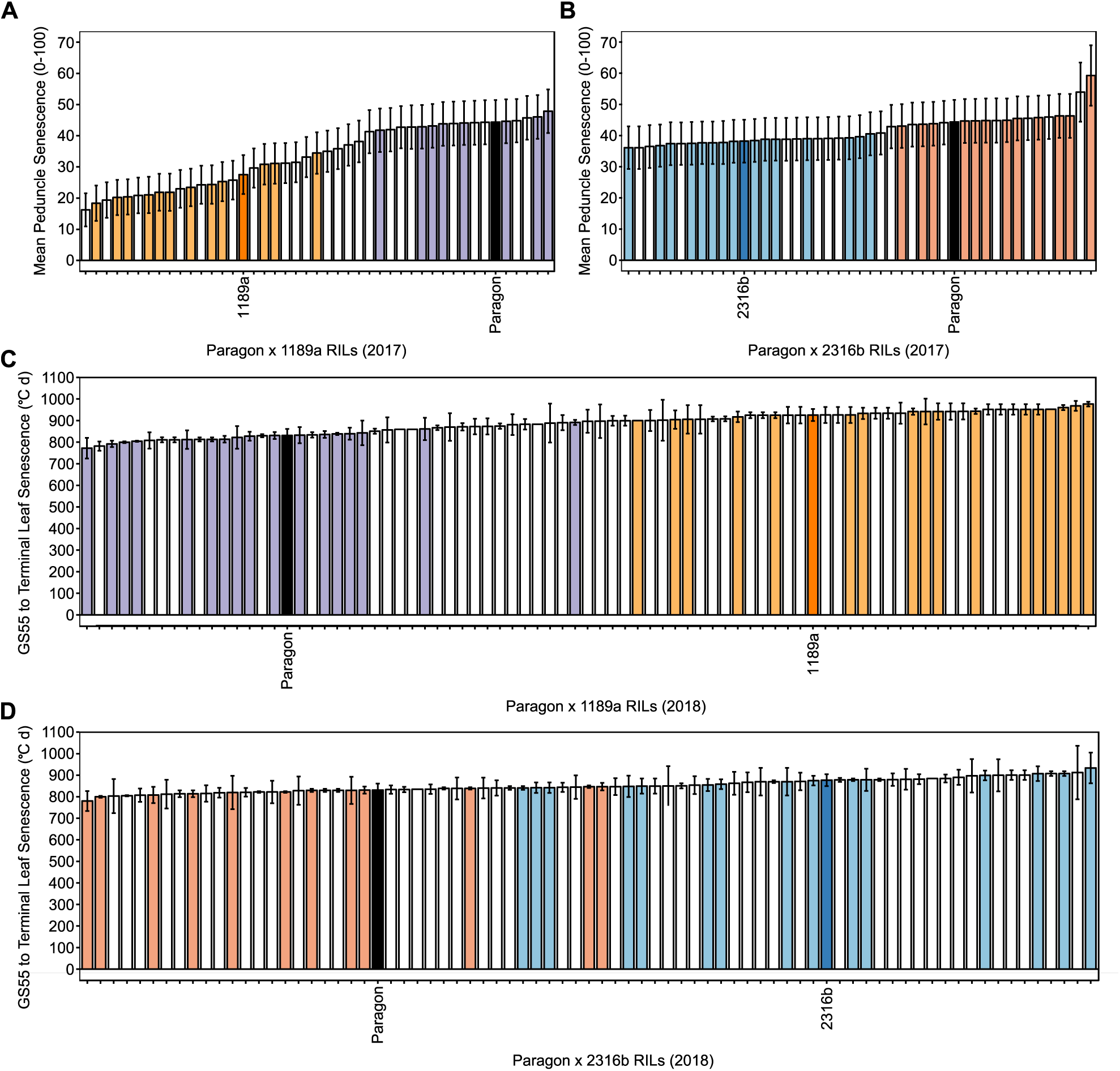
Senescence variation amongst F_4_ RIL populations, highlighting RILs included in bulks. Mean peduncle senescence scores for Paragon x 1189a **(A)** and Paragon x 231Gb **(B)** F_4_ Rlls grown in 2017. Duration of leaf senescence (from ear emergence) for Paragon x 1189a **(C)** and Paragon x 231Gb **(D)** F_4_ RILs grown in 2018. Coloured bars represent parents and RILs included in bulks, 1189a (orange), ‘staygreen’ (light orange, n = 17), ‘non-staygreen’ (light purple, n = 17); 231Gb (dark blue), ‘staygreen’ (light blue, n = 15), ‘non-staygreen’ (red, n = 12); Paragon (black). Classification and selection of RILs guided by multiple senescence metrics with intra- and inter-year comparisons performed.

### 1189a and 2316b Staygreen Traits are located on Chromosomes 6A and 6D

Following classification of F_4_ RILs into ‘staygreen’ and ‘non-staygreen’ types, DNA of selected individuals was pooled to form staygreen and non-staygreen bulks (Figure 3) and submitted for exome capture and sequencing. Reads were aligned to the *Triticum aestivum* cv. Chinese Spring RefSeq v1.0 (IWGSC *et al.*, 2018) and variants identified (Figure 4). Coverage of high confidence exons according to IWGSC gene annotation v1.1 (Alaux *et al.*, 2018; IWGSC *et al.*, 2018) ranged from ∼35 to 43 %, equating to over 131 million positions for each parental line or bulk. For each exome capture sample, a read depth ≥ 20 was recorded for 80 to 90 million positions, not exclusive to exons. Of these positions ∼57 K to 68 K were SNPs, with one third being characteristic EMS G:A or C:T transitions, with SNPs identified within the cv. Paragon sample being varietal (Supplemental Table S2).

**Figure 4.**
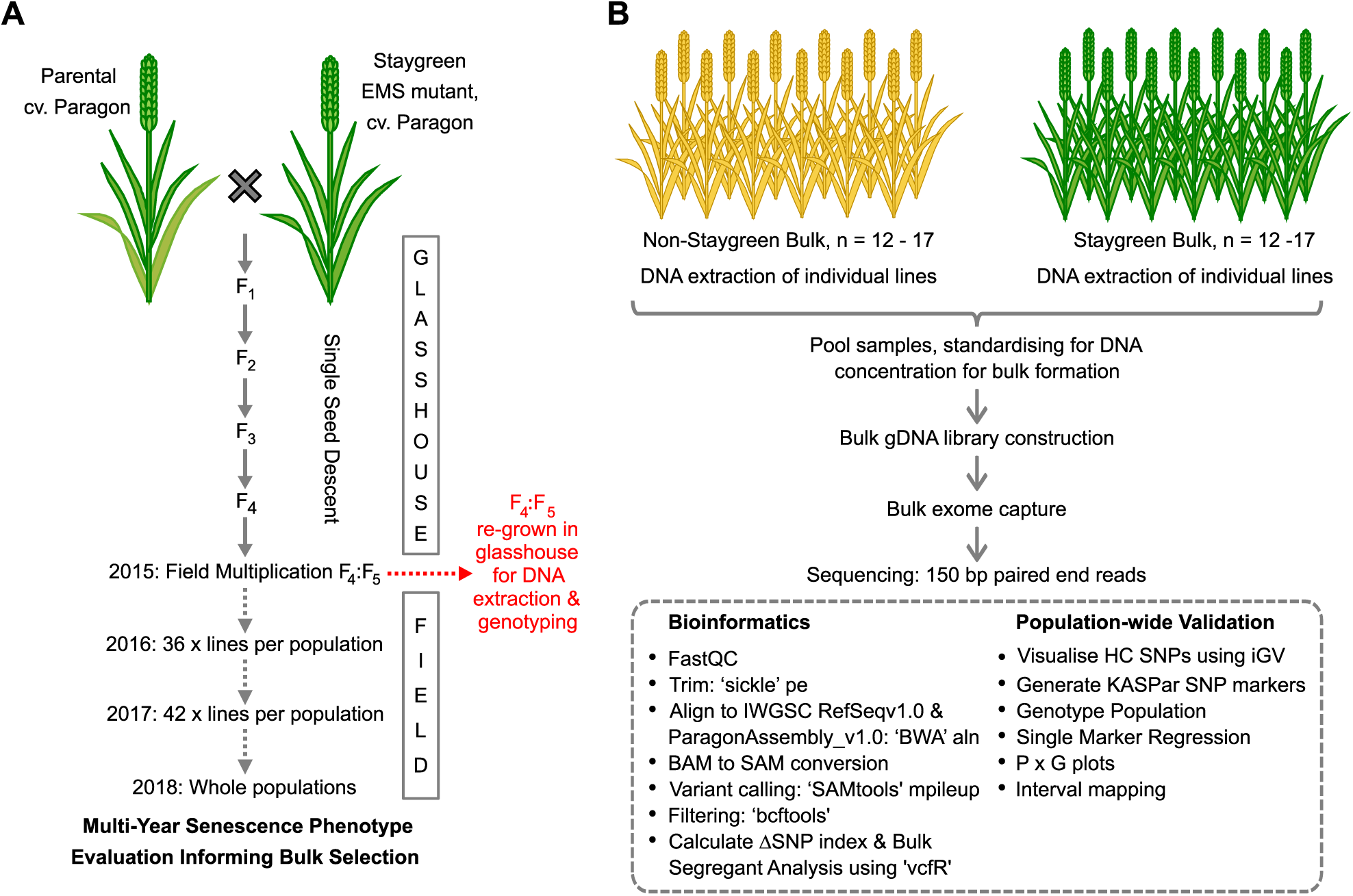
Development of RIL populations segregating for senescence traits and mapping strategy. **(A)** Staygreen mutants were crossed to cv. Paragon and F_4_ RIL population developed through SSD. Differing numbers of RILs underwent phenotypic assessment in the field between 2016-2018. **(B)** Following classification of RILs as ‘staygreen’ or ‘non-staygreen’, DNA of selected RILs was pooled to form contrasting bulks and submitted for exome capture and sequencing alongside parents. Sequencing reads were subject to QC analysis and processing prior to read mapping and variant detection facilitating performance of BSA. KASP markers were developed for SNPs of interest for genotyping of entire RIL populations and marker association analysis performed.

Mapping traits using BSA aims to identify variants that are both enriched and depleted between contrasting bulks across genetic region(s) (Michelmore and Kesseli, 1991). To standardise for variation in read depth a SNP index is calculated by dividing DV/DP (Number of reads for the alternative allele/Total read depth). SNPs associated with delayed senescence phenotypes should be present in most reads within ‘staygreen’ bulks, SNP index = 1, but absent in the ‘non-staygreen’ bulk, SNP index = 0 (ΔSNP index = 1). Variants for which SNP index = 1 across both bulks are varietal (ΔSNP index = 0).

Upon removal of varietal SNPs, 37 495 and 38 947 transition type SNPs (DP ≥ 5) were retained across the 1189a and 2316b bulks, respectively. Filtering SNPs based on SNP index (non-staygreen ≤ 0.05 and staygreen ≥ 0.95) identified independent regions on chromosomes 6A and 6D enriched for 1189a and 2316b alleles, respectively (Figure 5). A ΔSNP index = 1 was calculated for 6 SNPs located on chromosome 6A for 1189a (Table 2), and 10 SNPs located on chromosome 6D for 2316b (Table 3).

**Figure 5.**
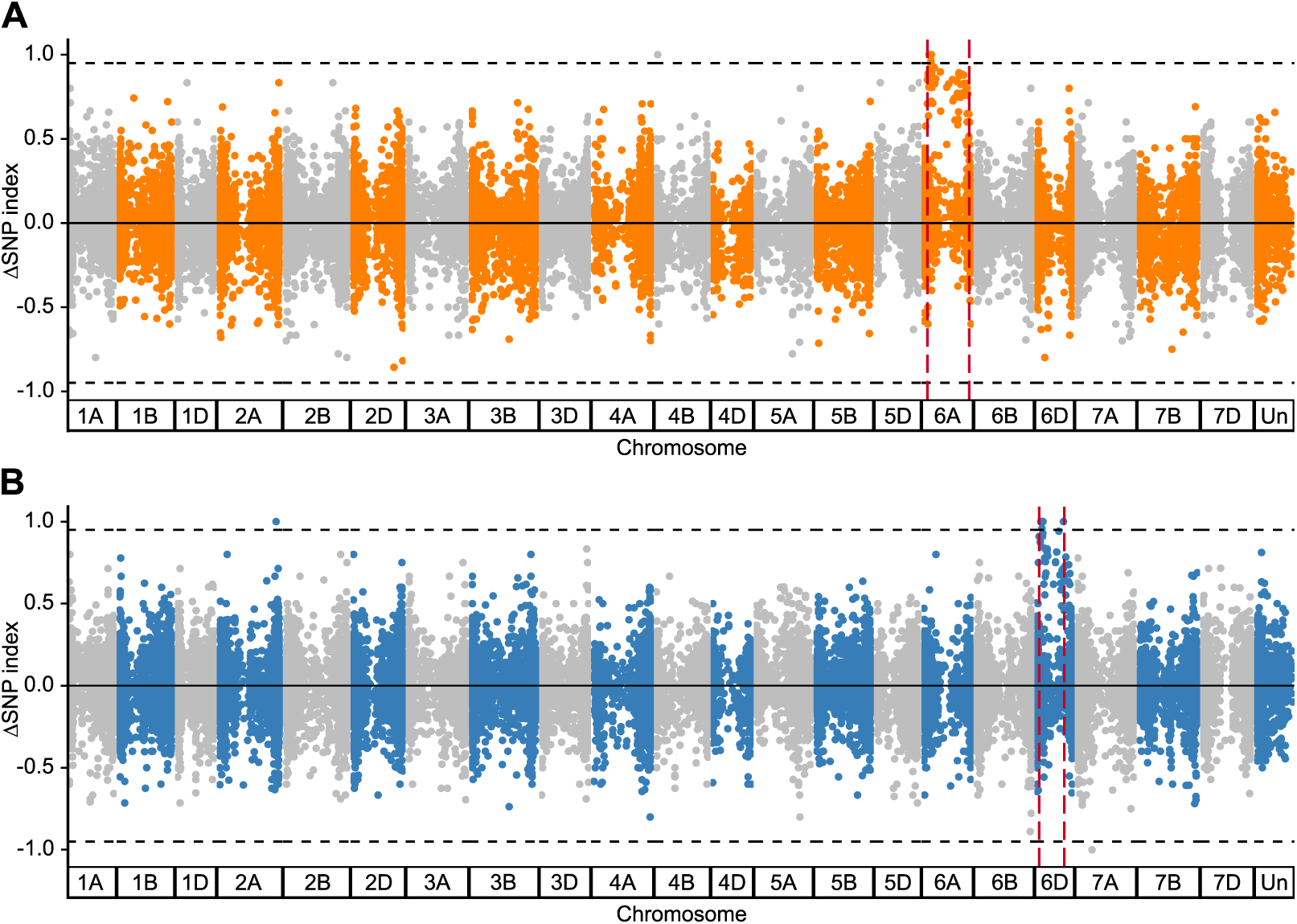
Bulk segregant analysis maps 1189a and 2316b staygreen traits to two independent loci. Calculation of SNP index {SNP index_sG_ - SNP index_non-sG_) identifies SNPs characteristic of EMS-mutagenesis are enriched across single regions of chromosome GA for 1189a **(A)** and chromosome 6B for 231Gb **(B)**. SNPs plotted are G:A and C:T transitions (1189a, n = 37 495; 231Gb, n = 38 947) against physical position of *Triticum aestivum* cv. Chinese-Spring RefSeq vl.1 (IWGSC *et al.*, 2018).

### 1189a and 2316b staygreen traits are underpinned by mutations in *NAM-1*

To determine if SNPs identified as enriched during BSA are causative variant effect prediction was conducted. For 1189a, 13 SNPs located on chromosome 6A are predicted to encode missense mutations, with 9 deleterious to protein function (SIFT ≤ 0.01) (Table 2). For 2316b, 5 SNPs located on chromosome 6D are predicted to encode missense mutations, 3 of which are deleterious (SIFT ≤ 0.02) (Table 3). These deleterious SNPs were prioritised as gene candidates according to ΔSNP indices. Results for 1189a and 2316b converge upon *NAM-A1* (Table 2) and *NAM-D1* (Table 3) (ΔSNP index = 1); homoeologues of known senescence regulator *NAM-B1* (Uauy *et al.*, 2006a; Uauy *et al.*, 2006b)

To validate these results, we developed KASP markers for SNPs located on chromosomes 6A (1189a) and 6D (2316b) (Supplemental Table S3, Supplemental Table S4). Genotyping of F_4_ RIL populations facilitated genetic map construction utilising all available RILs (Figure 6A-B). Single marker association analysis reported increasing phenotypic associations with proximity to *NAM-A1*, -log_10_(*P*) ≥ 3.5, of 1189a (Figure 3C) and *NAM-D1*, -log_10_(*P*) ≥ 2.35, of 2316b (Figure 3D) (Holm corrected for multiple testing), with size of association reflecting subtlety of senescence phenotype (Figure 1). Here, we mapped the 1189a interval to a 16.7 Mb region on chromosome 6A and the 2316b interval to a 4.8 Mb region on chromosome 6D, containing 142 and 56 high confidence genes, respectively (IWGSC *et al.*, 2018).

**Figure 6.**
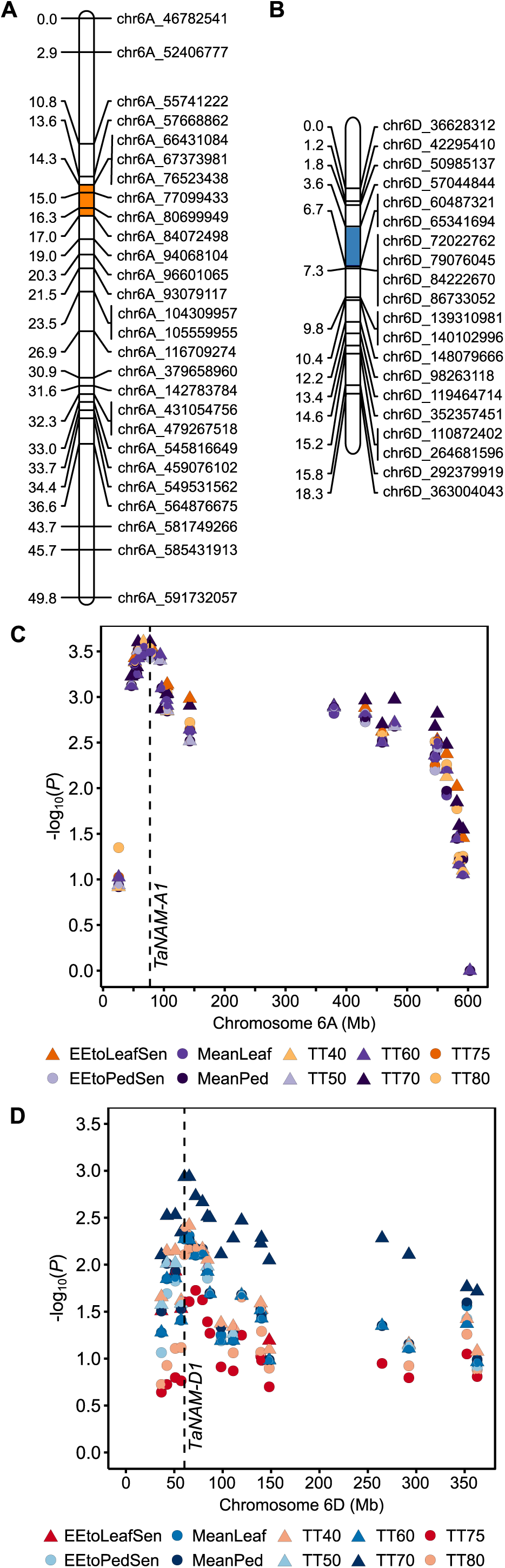
Genetic mapping of *NAM-1* homoeologues is supported by association analysis. KASP markers were developed for validation of positions enriched in staygreen bulks, F_4_ RILs genotyped, and genetic maps constructed for regions on chromosome 6A for 1189a **(A)** and 6D for 2316b **(B)**. Single marker association tests were performed using mapped SNPs and senescence phenotypes of Paragon x 1189a **(C)** and Paragon x 2316b **(D)** F_4_ RILs (n= 75) (2018, 2 replicates). Marker associations increase with proximity to *NAM-Al* and *NAM-01*, confirming candidate likelihood. Highlighted regions indicate 8SNP index= 1 and include SNPs within *NAM-Al* (chr6A_77099433) and *NAM-01* (chr6D_60487321). Markers named according to RefSeq vl.0 (IWGSC *et al.*, 2018). Marker distances calculated in MapDisto v2.0 (Heffelfinger *et al.*, 2017). Maps constructed using MapChart (Voorrips, 2002).

### 1189a and 2316b represent novel sources of *NAM-1* allelic variation

SNPs identified in *NAM-1* homoeologues encode missense mutations (Table 2, Table 3). Isoleucine replaces threonine at amino acid (AA) position 159 of NAM-A1 for 1189a (T159I). Glycine replaces glutamate at AA 151 of NAM-D1 for 2316b (G151E) (Figure 7). NAC transcription factors operate as heterodimers and homodimers, which ensures stable-DNA binding (Olsen *et al.*, 2005). The G151E and T159I AA substitutions are located within subdomain D of the NAC transcription factor domain at highly conserved positions involved in DNA-binding (Ooka *et al.*, 2003; Ernst *et al.*, 2004; Welner *et al.*, 2012; Harrington *et al.*, 2019a). Examination of the NAC domain crystal structure show variants mirror one another, with the T159I AA variant located one residue after the β4 structure, and G151E AA variant located one residue before the β5 structure of the antiparallel β-sheet secondary structure (Ernst *et al.*, 2004). The substitution of positively charged threonine for hydrophobic isoleucine, or glycine for a large, positively charged glutamate may affect NAM protein function by altering protein dimerization, as demonstrated for alternative EMS-induced mutations in NAM-A1 by Harrington *et al*. (2019a) (Figure 7).

**Figure 7.**
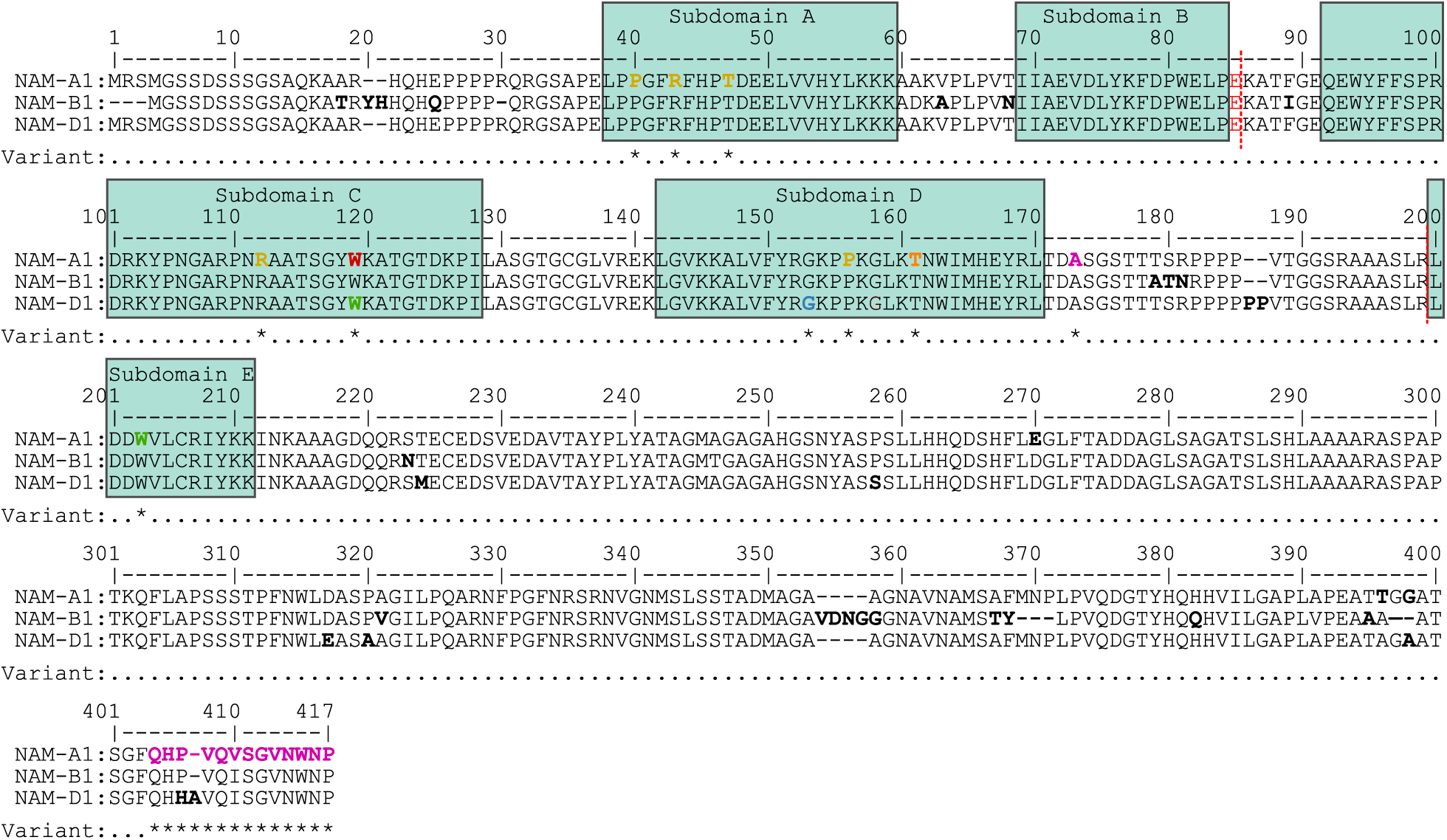
Sequence alignment of wheat NAM-1 homoeologous proteins. Boxes denote NAC subdomains (Ooka *et al.*, 2003). Bold black lettering indicates homoeologous variation, and red dashed lines exon-junctions. Asterisks and bold coloured lettering indicate known variants, including AA substitutions identified for 1189a in orange (G159E, NAM-Al), 2316b in blue (TlSl, NAM-D1), natural NAM-Al variants in pink (Cormier *et al.*, 2015), EMS-induced NAM-Al missense mutations in yellow (Harrington *et al.*, 2019a), EMS-induced knockout mutations in green (Avni *et al.*, 2014) and red (Pearce *et al.*, 2014). In *Triticum aestivum NAM-Bl* is largely non-functional due to a +1 bp frameshift mutation, or complete deletion (Uauy *et al.*, 2006a; Hagenblad *et al.*, 2012; Alhabbar *et al.*, 2018a; Alhabbar *et al.*, 2018b). Accession numbers are, NAM-Al, TraesCS6A02G108300.1; NAM-Bl, UniProtKB/Swiss-Prot: A0SPJ4.1 *(Triticum turgidum ssp. Dicoccoides);* NAM-D1, TraesCS6D02G096300.1.

### Differential Inheritance of 1189a and 2316b Staygreen Traits

RIL populations were developed by single seed descent (SSD) in the glasshouse with segregation of senescence phenotypes unobserved in earlier generations. Phenotype x Genotype plots constructed for Paragon x 1189a and Paragon x 2316b F_4_ RILs homozygous for alternative *NAM-1* alleles form distinct groups, *P* < 0.0001 (Figure 8, Supplemental Figure S6). Assessment of RILs heterozygous for the *NAM-A1* mutation reveal the allele is dominantly inherited, as RILs resemble those homozygous for the mutation, *P* > 0.3, not the cv. Paragon allele, *P* < 0.01 (Figure 8).

**Figure 8.**
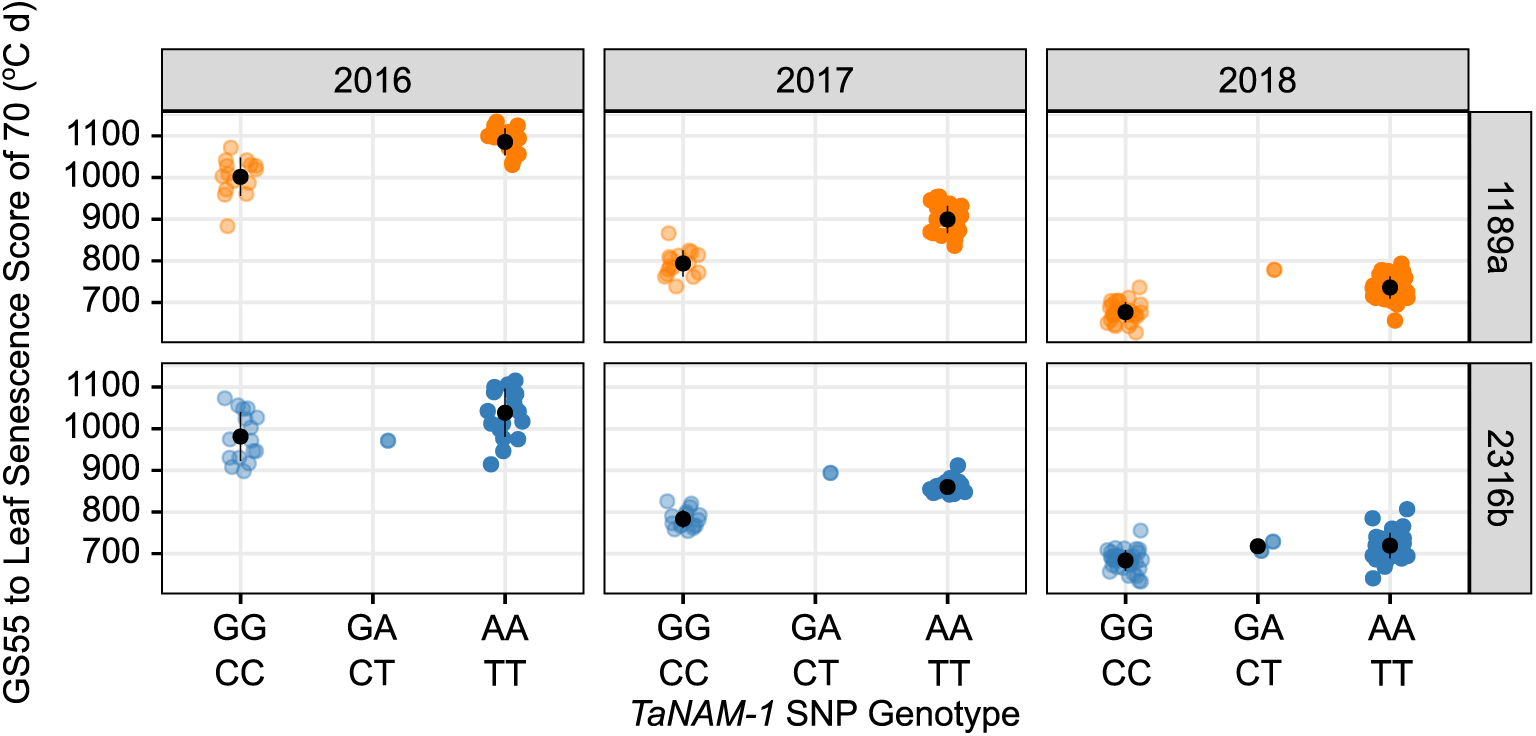
Phenotype x genotype plots illustrate staygreen traits are dominantly inherited. Scatterplot displaying TT70 values against *NAM-1* genotype of Paragon x 1189a (orange, top) and Paragon x 2316b (blue, bottom) F_4_ RI Ls in 2016 to 2018 (left to right). G/C cv. Paragon allele, A/T mutant allele. Phenotypic differences between contrasting homozygotes are significant, *P* < 0.001. Heterozygotes display intermediate phenotypes, *P* > 0.05. Mean phenotypic value, n = 1-3. Black circle, genotypic mean± SD. Differing numbers of RI Ls, including heterozygotes, grown each year, 2016, n = 36; 2017, n = 43; 2018, n > 75.

Inheritance of the 2316b *NAM-D1* mutation is more complex. In 2016 and 2018, senescence phenotypes of heterozygous Paragon x 2316b F_4_ RILs were indistinct from RILs homozygous for either *NAM-D1* allele (2016, *P* > 0.7; 2018, *P* > 0.28). In 2017 the *NAM-D1* mutation appears dominantly inherited, as senescence of RILs heterozygous and homozygous for the *NAM-D1* mutation was similarly delayed compared to RILs homozygous for the cv. Paragon allele, *P* < 0.01 (Figure 8). Together, this suggests the *NAM-D1* mutant allele is semi-dominant.

Validation of differential modes of inheritance for *NAM-A1* and *NAM-D1* mutations arose from phenotyping heterozygous RILs derived from independent crosses grown in Cambridgeshire and Norwich. Senescence profiles of homozygous RILs contrasting for *NAM-A1* alleles were significantly different, *P* < 0.01, with heterozygotes resembling RILs homozygous for the mutation, *P* > 0.66 (Supplemental Figure S7). Penetrance of the *NAM-D1* mutant allele is environmentally dependent. Differences in senescence phenotypes of homozygous RILs contrasting for *NAM-D1* alleles were significant in Cambridgeshire, *P* < 0.05, but not in Norwich, *P* > 0.5 (Supplemental Figure S7). *NAM-D1* heterozygotes more closely resemble RILs homozygous for the mutant allele, *P* < 0.6, compared to the cv. Paragon allele, *P* > 0.9, supporting semi-dominant inheritance of the *NAM-D1* mutation (Supplemental Figure S7).

## Discussion

### Two novel wheat senescence mutants which extend grain fill duration

Grain filling experiments confirmed the hypothesised positive relationship between staygreen traits and grain fill duration (Wiegand and Cuellar, 1981; Gelang *et al.*, 2000). Grain fill extensions reported for lines 1189a and 2316b mirror observed delays in onset of senescence (Figure 2, Supplemental Figure S2, Supplemental Figure S3). Differences in grain fill between staygreen lines and cv. Paragon occur towards the end of the rapid grain filling phase, which Neghliz *et al*. (2016) estimates to occur 39 daa for *Triticum aestivum* cv. Recital when grain moisture content reaches ∼45 %. Photosynthesis terminates halfway through this rapid phase, whereupon translocation of stored reserves and remobilisation of fructose and sucrose occurs (Takahashi *et al.*, 1993; Takahashi *et al.*, 2001). Alongside the significant differences in grain moisture content recorded (Supplemental Table 1), we propose a potential extension in the rapid grain filling phase of ∼5 days for 1189a and 2316b. The additional grain fill extension observed for 1189a likely relates to differences in the final lag phase of grain fill (Takahashi *et al.*, 2001), as unlike 2316b, grain moisture content was significantly greater compared to cv. Paragon on 23 ^rd^ July in both years, *P* < 0.001 (Supplemental Table S1). Delayed grain maturation of 1189a may disrupt depletion of stem reserves and deposition of triticin, glutenin and gliadin storage proteins occurring during this final phase (Takahashi *et al.*, 2001; Triboi *et al.*, 2003), potentially reducing grain quality and requires investigation.

Although the pattern of grain moisture decline for our staygreens is environmentally stable, dry grain weight accumulation is under greater environmental influence (Figure 2, Supplemental Figure S3). Extended grain fill duration of 2316b was not associated with increasing final grain weight, *P* > 0.05, but was for 1189a in 2018, *P* < 0.001, but not in 2017, *P* > 0.05 (Supplemental Table 1). In 2018, grain fill was curtailed by ∼4 to 5 days for all lines compared to 2017, with temperatures exceeding the 12-22 °C range considered optimal (Dias and Lidon, 2009; Farooq *et al.*, 2011). Therefore, the greater final grain weight of 1189a reveals an association between staygreen traits and stress tolerance, as reviewed by Gregersen *et al*. (2013) and Thomas and Ougham (2014).

Final grain weight and grain filling rate are significantly correlated (Dias and Lidon, 2009), with slower rates reducing remobilisation efficiency (Xie *et al.*, 2015). Grain filling rate of these staygreens may be affected, illustrated by the shallower gradients between timepoints when grain moisture contents are significantly different, and potentially slowest for 1189a (Figure 2, Supplemental Figure S3). The later termination of photosynthesis for 1189a, and greater availability of photosynthates, may counteract the reduction in stem reserve remobilisation under stress, explaining the perhaps contradictory final grain weight increase in 2018. Evidence of additional photosynthates sustaining grain fill of staygreens are the greater dry grain weights recorded for 1189a from 42 daa, and 2316b from 37 daa, *P* < 0.05 (Supplemental Table S1). Grain filling experiments by Borrill *et al*. (2015) support this, with flag leaves of *NAM-*RNAi lines producing 2 079 mg more glucose per plant compared to controls. For 2316b, these earlier increases did not improve final grain weight suggesting remobilisation efficiency may be impaired like 1189a, with earlier termination of senescence unable to compensate. Alternatively, any final grain weight improvement associated with 2316b may be diluted due to background mutations.

### *NAM-A1* and *NAM-D1* as gene candidates

Regions identified during BSA for 1189a and 2316b encode mutations in *NAM-1* homoeologues. *NAM-B1* is a known senescence regulator, previously identified in *Triticum turgidum ssp. dicoccoides* during dissection of locus *GPC-1* (Uauy *et al.*, 2006a; Uauy *et al.*, 2006b). In most hexaploid wheats *NAM-B1* is non-functional due to a +1 bp frameshift-encoding insertion or its complete deletion (Uauy *et al.*, 2006a; Hagenblad *et al.*, 2012; Lundström *et al.*, 2017). The role of *NAM-A1* and *NAM-D1* homoeologues in senescence was confirmed by Avni *et al*. (2014), with natural variation limited to A and B homoeologues with no D variants reported. Of the *NAM-A1* alleles detected one encodes an alanine to valine AA substitution between subdomains D and E (A171V), the other a frameshift-induced truncation mutation (Figure 7) (Cormier *et al.*, 2015).

SNPs within *NAM-1* homoeologues of 1189a and 2316b are considered deleterious to protein function (SIFT = 0) (Table 2, Table 3) and are completely associated with senescence phenotypes (Figure 6). Supporting the proposal of these mutations as causative is a reverse genetic study concerning *gpc-1* (*NAM-1*) TILLING mutants by Avni *et al*. (2014) (Figure 7). A delay in onset of senescence of ∼6 days and ∼3 days was reported for the *Triticum aestivum* cv. Express *gpc-a1* W196* truncation mutant and *gpc-d1* W114* knockout mutant, respectively (Avni *et al.*, 2014), matching the phenotypes recorded for 1189a and 2316b (Table 1). Likewise, senescence of *gpc-1* mutants and 1189a and 2316b progresses in parallel, terminating 6-10 days and 5 days later compared to cv. Express, respectively (Avni *et al.*, 2014; Supplemental Figure S2).

Within a protein-context, mutations within *NAM-A1* and *NAM-D1* are self-validating, encoding AA substitutions within subdomain D of the NAC domain (Figure 7) within an identified DNA binding region (Ooka *et al.*, 2003; Welner *et al.*, 2012). Mutations in subdomain D can alter NAM protein functionality, with the affected G151 and T159 residues conserved in over 65 % of NAC transcription factor encoding genes (Ooka *et al.*, 2003; Puranik *et al.*, 2012; Welner *et al.*, 2012; Fan *et al.*, 2014). In *Triticum turgidum* TILLING mutants, a P154L mutation in NAC subdomain D of NAM-A1 disrupted protein dimerization in absence of a senescence phenotype (Harrington *et al.*, 2019a), whilst a G133D mutation in subdomain D of NAM-A2, the *NAM-A1* paralog, significantly delayed peduncle senescence (Borrill *et al.*, 2019). Similarly, *NAM-1* homologs encoding subdomain D allelic variants in *NAM-G1* of *Triticum timopheevi* and *NAM-1* of *Hordeum vulgare* are associated with reducing grain protein content, *P* < 0.05, illustrating loss of function (Jamar *et al.*, 2010; Hu *et al.*, 2013).

Welner *et al*. (2012) propose one NAC monomer initially sub-optimally binds DNA with the other scanning and searching for a binding site. Changes in charge or polarity introduced by G151E and T159I mutations may disrupt initial DNA-binding to prevent dimerization, with NAM-1 homoeologues affected similarly due to structural palindromicity of residues (Ernst *et al.*, 2004; Welner *et al.*, 2012). Performance of Yeast-2-Hybrid and cell death assays for mutated NAM-1 proteins would test for altered binding activity, as performed by Harrington *et al*. (2019a), to identify residues critical to NAM-A1 protein function. The extremity of the 1189a staygreen phenotype compared to 2316b likely reflects homoeologous dominance of *NAM-A1* over *NAM-D1* and not mutation type, which phenotypic characterisation of *gpc-1* mutants supports (Avni *et al.*, 2014). However, involvement of linked mutations within mapped intervals not captured by exome sequencing, including promoter variants, cannot be disregarded, but together these data suggest *NAM-A1* and *NAM-D1* mutations are causative.

### Grain fill phenotypes of 1189a and 2316b compare favourably with NAM-1 variants

When characterising *GPC-B1* (*NAM-B1*) Uauy *et al*. (2006b) reported an association between the non-functional allele and longer grain filling period. Validating the association between variation and grain fill extension observed for 1189a (*NAM-A1*) and 2316b (*NAM-*D1) mutants are the results of Avni *et al*. (2014). From 42 daa spike moisture content of *gpc-1* mutants was greater compared to parental controls, *P* < 0.05, and remained so until 49 daa and 57 daa for the *gpc-d1* and *gpc-a1* mutant, respectively, matching the pattern of grain moisture loss recorded for 1189a and 2316b (Figure 2, Supplemental Figure S3). *NAM-A1* variants are common in Australian wheat cultivars with variation characteristic of mid to mid-late maturity types (Alhabbar *et al.*, 2018a), as observed for 1189a (Figure 2).

Contrary to the hypotheses of Wiegand and Cuellar (1981), Gelang *et al*. (2000) and Bogard *et al*. (2011) delayed senescence and grain fill extension, as associated with *NAM-1* variation, may not improve final grain weight. During grain fill, dry grain weights recorded for 1189a and 2316b were greater, *P* < 0.05 (Supplemental Table S1), contradicting the lower weights recorded for *NAM-1* RNAi lines, *P* < 0.05 (Borrill *et al.*, 2015) and similar dry spike weights recorded for *gpc-1* mutants, *P* > 0.05 (Avni *et al.*, 2014). Differences in final grain weight between *NAM-1* RNAi, 2316b and *gpc-1* mutant lines and controls were not significant, *P* > 0.05 (Avni *et al.*, 2014; Borrill *et al.*, 2015), however grain weight of 1189a was greater in 2018, *P* < 0.001 (Supplemental Table 1).

The influence of *NAM-1* variation on final grain weight depends on genetic background. Differences in TGW of isogenic lines carrying a non-functional *GPC-B1* copy were inconsistent and both higher and lower, *P* < 0.05 (Uauy *et al.*, 2006b), and likewise for cultivars carrying *NAM-A1* variants (Cormier *et al.*, 2015; Alhabbar *et al.*, 2018a). Borrill *et al*. (2015) hypothesises similarity in TGW of *NAM-1* RNAi lines and controls, *P* = 0.25, was due to inadequate starch synthase activity. Instead of contributing to grain filling activities, Borrill *et al*. (2015) demonstrated the additional sucrose synthesised by *NAM-1* RNAi lines was retained as stem fructan within the internodes. Improved stem fructan remobilisation is associated with TGW improvement (Zhang *et al.*, 2015), with fructan providing a source of water-soluble carbohydrates (WSCs) for sustainment of grain filling (Fischer, 2011; Borrill *et al.*, 2015). Greater final grain weight of 1189a in 2018, and recorded during grain filling for 2316b, *P* < 0.05 (Supplemental Table S1) demonstrate the novel *NAM-1* alleles we identified could increase TGW. Therefore, combining these *NAM-1* alleles with a mutation in gene *1-FEH-w3* regulating stem fructan remobilisation (Zhang *et al.*, 2015) could overcome problems associated with fructan retention (Borrill *et al.*, 2015) and consistently improve TGW.

### Dominance of NAM genes in regulation of wheat senescence

Genetic mapping of 1189a and 2316b converged upon homoeologous copies of known senescence regulator *NAM-B1* (Uauy *et al.*, 2006a). Similarly, a forward genetic screen of a *Triticum turgidum* cv. Kronos TILLING population to identify senescence mutants converged upon *NAM-A1* mutations (Harrington *et al.*, 2019b). The differential onset of senescence observed for 1189a and 2316b (Table 1) reflects the reported dominance of *NAM-1* homoeologues (Avni *et al.*, 2014), with our results the first forward genetic screen identifying *NAM-D1*.

*NAM-1* is a positive regulator of senescence, with expression upregulated following anthesis (Uauy *et al.*, 2006a) and is associated with transcriptional reprogramming. At 12 daa RNA-seq studies of *gpc-1* (*NAM-1*) mutants identified ≥ 691 differentially expressed genes, with protein catabolism and stress response genes upregulated, and photosynthetic and housekeeping genes downregulated (Cantu *et al.*, 2011; Pearce *et al.*, 2014; Borrill *et al.*, 2019). The role of *NAM* genes in senescence regulation is complicated by the *NAM-1* paralog, *NAM-2*. RNA-seq studies of tetraploid wheat *gpc-1* and *gpc-2* mutants identified *NAM-1* as dominant over *NAM-2*, with expression associated with 64 % of senescence regulated genes compared to 37 %, respectively (Pearce *et al.*, 2014). Phenotypic characterisation of *NAM-B2* mutants confirm its role in senescence regulation, however phenotypic differences were only significant when combined with mutations in *NAM-A1* or *NAM-A2, P* < 0.05 (Pearce *et al.*, 2014; Borrill *et al.*, 2019). Together, these findings illustrate the problems associated with identifying novel senescence regulators using forward genetic techniques.

Mutations identified in *NAM-1* homoeologues were penetrative and dominant (Figure 8), with their detection relatively unconfounded by homoeologues as cv. Paragon encodes a non-functional copy of *NAM-B1*. Mutations in genes acting downstream of *NAM* may affect expression of fewer genes resulting in subtler phenotypes. Inconsistencies between leaf and peduncle phenotypes (Borrill *et al.*, 2019; Harrington *et al.*, 2019a; Harrington *et al.*, 2019b), spatial variation or effects of background mutations may prevent identification of these. Differential senescence phenotypes were confirmed for multiple *Triticum aestivum* cv. Paragon EMS mutants, with resource worth further exploration, including sequencing of *NAM-1* and *NAM-2* homoeologues of staygreen mutants to identify causative mutations due to their known role in senescence (Avni *et al.*, 2014; Pearce *et al.*, 2014; Borrill *et al.*, 2019; Harrington *et al.*, 2019a). Conversely, early senescing mutants may encode gain of function mutations affecting *NAM-1* gene regulatory targets including C2C2-CO like transcription factors, RWD-RK or GRAS genes identified during transcriptional network modelling (Borrill *et al.*, 2019; Harrington *et al.*, 2019c).

## Conclusions

Here, we confirm the central role of *NAM-1* in the genetic regulation of wheat senescence through identifying novel mutant *NAM-A1* and *NAM-D1* alleles through a forward genetic screen. Both mutations occurring within subdomain D of the NAC domain, therefore highlighting the importance of this protein in modulating NAM-1 function. Altered senescence profiles associated with these mutations are independent of heading-date variation and contribute to a grain fill extension and potential increased grain weight.

## Materials & Methods

### Plant Material

#### Mutagenesis and Initial Screen

7000 seeds of spring wheat cv. Paragon were treated with a 1 % EMS solution for 16 hours to obtain a 50 % lethal dose for viability, according to Rakszegi *et al*. (2010). 3500 M_1_ seeds were sown to obtain M_1_ plants, with the surviving 3461 bagged to ensure self-fertilisation. Two M_2_ seeds were sown per line, deriving ‘a’ and ‘b’ sibling lines (n = ∼6922) and advanced via multiple rounds of self-fertilisation to the M_5_ generation. In spring 2006 ∼6500 M_5_ lines were grown as single ear rows in 1 m^2^ plots at Church Farm, Bawburgh (52°38’N 1°10’E) (JIC). A single visual assessment of the 6500 M_5:6-_ lines identified 18 early senescing and 43 staygreen mutants. Original data available from: www.wgin.org.uk/wgin_2003-2008/index.php?area=Resources&page=results. In 2007, forward genetic screening of the cv. Paragon EMS population was repeated under low nitrogen conditions (100 kg N ha-^1^). Seed of the M_5:6_ generation (n = 6500) was sown as single rows in 1 m^2^ plots. Phenotypic observations identified, or re-confirmed, differential senescence of ∼80 lines (data not shown). In 2008, 70 differentially senescing lines were included in a Nitrogen Use Efficiency screen and grown as replicated 1 m^2^ plots receiving 20 kg N ha^-1^ and 240 kg N ha^-1^ (n = 3). 54 lines were subject to glasshouse experimentation, with lines 555a, 862a, 1389a, 2056a and 2514a undergoing in-depth physiological characterisation by Derkx *et al*. (2012). This study concerns genetic mapping of 1189a and 2316b, chosen based on their environmentally stable, differential, staygreen phenotypes unconfounded by heading variation, and strong agronomic performance.

#### RIL Population Development

RIL populations segregating for senescence traits were developed for trait mapping purposes. 1189a and 2316b were crossed to parental *Triticum aestivum* cv. Paragon, and F_4_ populations developed through single seed descent (SSD); Paragon x 1189a (n = 85), Paragon x 2316b (n = 95). Field multiplication of F_4_ seed was conducted in Summer 2015, with RILs sown in numerical order in 1 m^2^ plots (3-rows, 40 cm spacing) in October 2014 (Figure 4A).

#### Field Experimentation

In-field phenotyping of F_4_ RIL populations was conducted between 2016 and 2018. Experiments were performed at Church Farm, Bawburgh (52°38’N 1°10’E), JIC. In 2016, 36 RILs per population were sown as unreplicated 1 m^2^ spaced plots on 26/10/2015. In 2017 and 2018 experiments consisted of replicated 6 m^2^ plots incorporating additional RILs per population (2017, n = 43, 3 replicates; 2018, n ≥ 75, 2 replicates) and were sown on 26/10/2016 and 12/10/2017 (Figure 4A). All experiments followed a randomised complete block design, with control plots (cv. Paragon, Soissons, 1189a and 2316b mutant lines) randomly sown throughout. Experimental seed was produced during multiplication of RILs in 2015 when the most recent source or resulted from the preceding year.

The soil at Church Farm is sandy loam overlying alluvial clay. Experiments were rainfed but required supplemental irrigation in 2017. Seeds were dressed with Redigo Dieter (Bayer CropScience, Germany) and sown at a rate of 2750 seeds per 6 m^2^ (∼300 plants m^-2^). Fertiliser was applied over three occasions from late February to the end of April, totalling 214 to 228.5 kg N ha^-1^ and 62 kg SO_3_ ha^-1^. Plots received standard fungicide and herbicide treatment.

#### Phenotypic Assessment

Phenotyping was conducted at the plot level. Ear emergence (GS55) was scored when 50 % of ears had emerged halfway from the flag leaf (Zadoks, *et al.*, 1974). Senescence was scored visually every 2 to 4-days from anthesis using a 0 to 100 scale (intervals of 5). Flag leaves were scored according to the proportion of leaf yellowing. A score of 5 represents leaf tip necrosis, whilst 100 indicates complete chlorosis or death (Pask *et al.*, 2012). Multiple flag leaves were assessed simultaneously to give a plot score. Peduncle senescence was scored in 2017 and 2018 and assessed as the percentage of yellow peduncles (top 3-5 cm) per plot based for 3 to 4 batches of 10 tillers.

To interpret senescence dynamics senescence scores were plotted against thermal time in day °C from ear emergence (T_0_) to standardise for heading variation. For thermal time calculation mean daily temperatures were calculated using minimum and maximum daily temperatures recorded by Church Farm weather station (Location: 52°37’ 52.29” N, 1°10’ 23.57” E). Using time-course senescence data corresponding to RILs and controls, senescence profiles were quantified and RILs classified by deriving senescence metrics. Senescence metrics include mean senescence, onset, duration (from ear emergence or onset to terminal senescence) and thermal time to different leaf senescence scores. Onset and termination of senescence were considered the first time points recording senescence scores above 10 or 90, respectively. Calculation of time taken (in day °C) to reach specific senescence scores (TT30, TT40…) is similar to MidS (50 % senescence) (Christopher *et al.*, 2016), with senescence assumed to progress linearly between scoring points and time interpolated.

#### Grain Filling Experiments

To explore the relationship between senescence and grain filling traits, in 2017 and 2018 grain weight and moisture content were recorded for 1189a, 2316b and cv. Paragon from anthesis to maturation. To standardise for developmental differences between tillers, ∼50 ears per plot per genotype (2017, n = 1; 2018, n = 2) were tagged when 1 to 2 cm of peduncle became exposed. At 4 to 5-day intervals, starting from anthesis, 5 tagged ears per plot were sampled and sealed into a labelled ziplock bag to prevent moisture loss. Ears were then refrigerated whilst awaiting dissection (maximum 10 hours post sampling) and senescence of sampled plots scored. 10 grains per ear were dissected from the central region of the spike from positions 1 and 3 (the oldest grains) and placed into a single Eppendorf tube and weighed to determine fresh grain weight. With lids open, tubes were transferred to a 65 °C drying oven for grains to dry down, leaving the oven door ajar to prevent condensation. After 48 hours tubes were re-weighed to determine final dry grain weight and grain moisture content (%) calculated by dividing dry by fresh grain weight.

#### DNA Extraction

Seeds of Paragon x 1189a and Paragon x 2316b F_4_ RIL populations, cv. Paragon, 1189a and 2316b mutant parents were sown into individual cells of 96-well seed trays containing peat and sand and transferred to the glasshouse following 2-3 days of cold treatment. When plants reached the 3-leaf stage 5 cm of leaf tissue was harvested, concertinaed, and collected into a 96-well collection tray (Qiagen, 19560) and material stored at -80 °C until required. DNA was extracted using the Qiacube® (Qiagen, Germany) according to the QIAamp 96 DNA Qiacube HT Kit Protocol. DNA quality and quantity were analysed using a DS-11 Spectrophotometer (Denovix, DE, USA), Qubit-4 Fluorometer (dsDNA BR assay, Q32850, Thermofisher) and by running a DNA sample on an agarose gel (1 %) to detect high molecular weight DNA.

#### Exome Capture

RILs were classified as ‘staygreen’ or ‘non-staygreen’ based on similarity of phenotype to parental lines (1189a or 2316b and cv. Paragon) using calculated senescence metrics. RILs for inclusion in bulks were selected based on within and between-year concordance of senescence metrics. Details concerning bulk selection are found in Supplementary Material 2. DNA of selected RILs was pooled, standardising for DNA concentration to ensure equal RIL representation (1189a, n = 17 for both bulks; 2316b, n = 15 for staygreen, n = 12 for non-staygreen) (Figure 4B). Quality and quantity of bulked DNA was checked by running a DNA sample on an agarose gel (1%) and final DNA concentration determined using the Quibit-4 Fluorometer (dsDNA BR assay, Q32850, Thermofisher).

Exome capture was used to sequence predicted gene-encoding regions (Krasileva *et al.*, 2017) of the four bulks and three parents (1189a, 2316b and cv. Paragon). Library preparation, amplification and sequencing was performed by Novogene (Hong Kong) using the SeqCapEZ probe set 140430_Wheat_TGAC_D14_REZ_HX1 for *Triticum aestivum* (Roche, Nimblegen, WI, USA), described by Krasileva *et al*. (2017). Libraries were sequenced using the HiSeq4000 platform (illumina, CA, USA) producing 150 bp paired end reads. Sequencing read quality was analysed using FastQC (Andrews, 2014) with low quality sequences and adapter remnants removed using ‘sickle’ (version 1.2; paired end (pe) mode, default parameters (-q 20 -l 15) (Joshi and Fass, 2011).

#### Sequence Alignment & Bulk Segregant Analysis

Processed reads were aligned to *Triticum aestivum* cv. Chinese Spring RefSeq v1.0 (IWGSC *et al.*, 2018) using ‘BWA’ (version 0.7.17; command aln, default parameters (-n 4); sampe, default parameters (-n 10 -N 0)) (Li and Durbin, 2009). Aligned read pairs were retained and files converted from Binary Alignment/Map (BAM) to Sequence Alignment/Map (SAM) format, ordered, duplicates removed and indexed using ‘samtools’ (version 1.7; command view -f2, -S -h -u -b -o; sort -o; rmdup; index) (Li *et al.*, 2009). Variant calling was performed simultaneously for contrasting bulks and parents using ‘samtools’ (version 1.7; command mpileup, default parameters (-g -t DP, DV, DPR, INFO/DPR), producing a labelled output retaining read depth and quality information. Output files were parsed using bcftools (version 1.8) (Li and Barrett, 2011) using the multi-allelic calling mode (-mv) (Danecek *et al.*, 2014) generating multi-sample Variant Call Format (.vcf) files. Variants assigned a quality score (“QUAL”) above 20 were retained to improve accuracy, and positions with missing data removed to aid genotypic comparison using bcftools (version 1.8; command filter -I “%QUAL > 20” -e “FORMAT/GT[0-1] = “./.”.) (Li and Barrett, 2011).

To conduct BSA multi-sample vcf files corresponding to each staygreen were imported into RStudio (RStudio team, 2015), R version 3.5.2 (R Core Team, 2018), using package ‘vcfR’ (Knaus and Grünwald, 2017). To identify variants enriched in staygreen versus non-staygreen bulks SNP indices were calculated for each position by dividing DV (the number of reads with the alternate allele) by DP (total read depth). Variants for which SNP index = 1 across bulks are varietal, occurring between cv. Paragon and Chinese-Spring, were removed. Remaining variants were filtered based on SNP index (staygreen > 0.9, non-staygreen < 0.1) and depth (DP > 3). Mutations characteristic of EMS mutagenesis (G:A and C:T transitions) were prioritised for investigation and visualisation in iGV (Integrated Genomics Viewer) (Robinson *et al.*, 2011). Variant enrichment across the genome was visualised by plotting ΔSNP index (SNP index_SG bulk_ – SNP_NSG bulk_) using ggplot2 (Wickham *et al.*, 2018).

#### Genotyping and Genetic Mapping

For SNPs of interest homoeologue-specific KASP markers were designed and RIL populations genotyped. Primer assay mixes contained 46 μl dH_2_O, 30 μl common primer (100 μM) and 12 μl of each tailed primer (100 μM). Genotyping assays were performed in 384-well format with a 2.5 μl KASP reaction volume consisting 14-18 ng DNA, 1.25 μl PACE 2 x Mastermix (3crbio, UK), 1.25 μl dH_2_0 and 0.047 μl primer assay mix. KASP assays were performed using a hydrocycler or thermocycler, with cycling conditions as follows: 15 mins at 94 °C, 10 cycles of 20 s at 94 °C, 60 s at 65-57 °C (decreasing by 0.8 °C per cycle), followed by 26-40 cycles of 20 s at 94 °C, 60 s at 57 °C. PCR cycling conditions required optimisation for some markers. KASP markers developed are listed in Supplemental Table S3 (1189a) and Supplemental Table S4 (2316b). Fluorescence was measured using a Pherastar plate reader (BMG Labtech, Germany) and data analysed using KlusterCaller software (version 4.1; LGC Genomics, UK).

Genetic maps for RIL populations were constructed from KASP genotyping data using the ‘Kosambi’ mapping function, specifying a LOD score of 6 in MapDisto v2.0 (Heffelfinger *et al.*, 2017). Loci were initially ordered using ‘Automap’ and ‘Find group’ functions and loci with missing data removed. Loci were reordered using ‘ripple order’, ‘check inversions’ and ‘order sequence’ functions with optimal order chosen based on shortest computed length and greatest increase in LOD score. Genetic maps were drawn using MapChart (Voorrips, 2002).

Single marker analysis was performed to identify peak marker-trait associations using package ‘AssocTests’ (Wang *et al.*, 2017). To correct for multiple testing outputted P-values were adjusted using the Holm correction. To determine if identified SNPs were directly causative variant effect prediction (McLaren *et al.*, 2016) was performed and SIFT score outputted (0 = deleterious, 1 = tolerable). To confirm the mendelian mode of inheritance of staygreen traits phenotype x genotype plots were constructed for peak markers using package ‘r/qtl’ (Arends *et al.*, 2010).

## Data Analysis

Data analysis was performed using R (version 3.5.2) (R Core Team, 2018) in RStudio (RStudio team, 2015) and data manipulated using packages ‘data.table’ (Dowle *et al.*, 2019), ‘dplyr’ (Wickham *et al.*, 2018), ‘plyr’ (Wickham, 2015) and ‘tidyr’ (Wickham *et al.*, 2019). Senescence metrics were derived from raw senescence phenotyping data in absence of spatial correction and means calculated per line when replicated. Analysis of senescence and grain filling profiles of individual lines or groups was conducted using linear mixed modelling involving packages ‘lme4’ (Bates *et al.*, 2019) and ‘lmerTest’ (Kuznetsova *et al.*, 2017). Tukey-Post hoc tests were performed using package ‘lsmeans’ (Lenth, 2018). Graphs were constructed using ‘ggplot2’ (Wickham *et al.*, 2018).

## Accession numbers

Raw exome capture sequencing reads for all samples have been deposited on the European Nucleotide Archive (ENA) (PRJEB40428).

## Supplementary Material 1

**Supplemental Figure S1** Senescence phenotypes for lines 1189a & 2316b (2016 & 2018)

**Supplemental Figure S2** Senescence phenotypes corresponding to grain filling experiments (2018)

**Supplemental Figure S3** Senescence and grain filling phenotypes (2017)

**Supplemental Figure S4** TT70 scores for Paragon x 1189a F_4_ RILs (2016 to 2018)

**Supplemental Figure S5** TT70 scores for Paragon x 2316b F_4_ RILs (2016 to 2018).

**Supplemental Figure S6** Additional Phenotype x Genotype plots illustrating mode of inheritance.

**Supplemental Figure S7** Independent confirmation of modes inheritance for *NAM-1* mutations.

**Supplemental Table 1** KASP primers for 1189a

**Supplemental Table 2** KASP primers for 2316b

**Supplemental Table 3** Differences in grain filling parameters, pairwise-comparison (2017 & 2018)

**Supplemental Table 4** Exome Capture Coverage and identified SNPs

**Supplementary Material 2** RIL senescence classification & bulk selection

## Acknowledgments

The authors wish to thank the JIC field experimentation team, without whom field trials would not have been possible, in addition to horticultural services and glasshouse staff. Thanks go to Burkhard Steuernagel for bioinformatics assistance, members of the Uauy group (JIC) for idea exchange, Luzie Wingen for statistical support, Rajani Awal and Richard Goram for tissue collection and DNA extraction. Thank you to Clare Lister for editorial assistance. The *Triticum aestivum* cv. Paragon population was developed by Robert Koebner and Leodie Alibert.

