## Supplementary Material 1 for "Delaying or Delivering: Identification of novel NAM-1 alleles which delay senescence to extend grain fill duration of wheat"

**Figure S1. Confirming environmental stability of senescence phenotypes for lines 1189a & 2316b.** Onset of leaf senescence was delayed for both mutants in 2016 **(A)**, alongside peduncle senescence in 2018 **(B)**. Differences in senescence progression were usually significant, P-value < 0.1 (Table 1). Senescence was scored visually using a 0-100 scale every 2-4 times per week from ear emergence (GS55), with scoring dates converted to thermal time (day °C). Mean ± SEM, n ≥ 4. Lines shown, cv. Paragon (black), 1189a (orange), 2316b (blue).

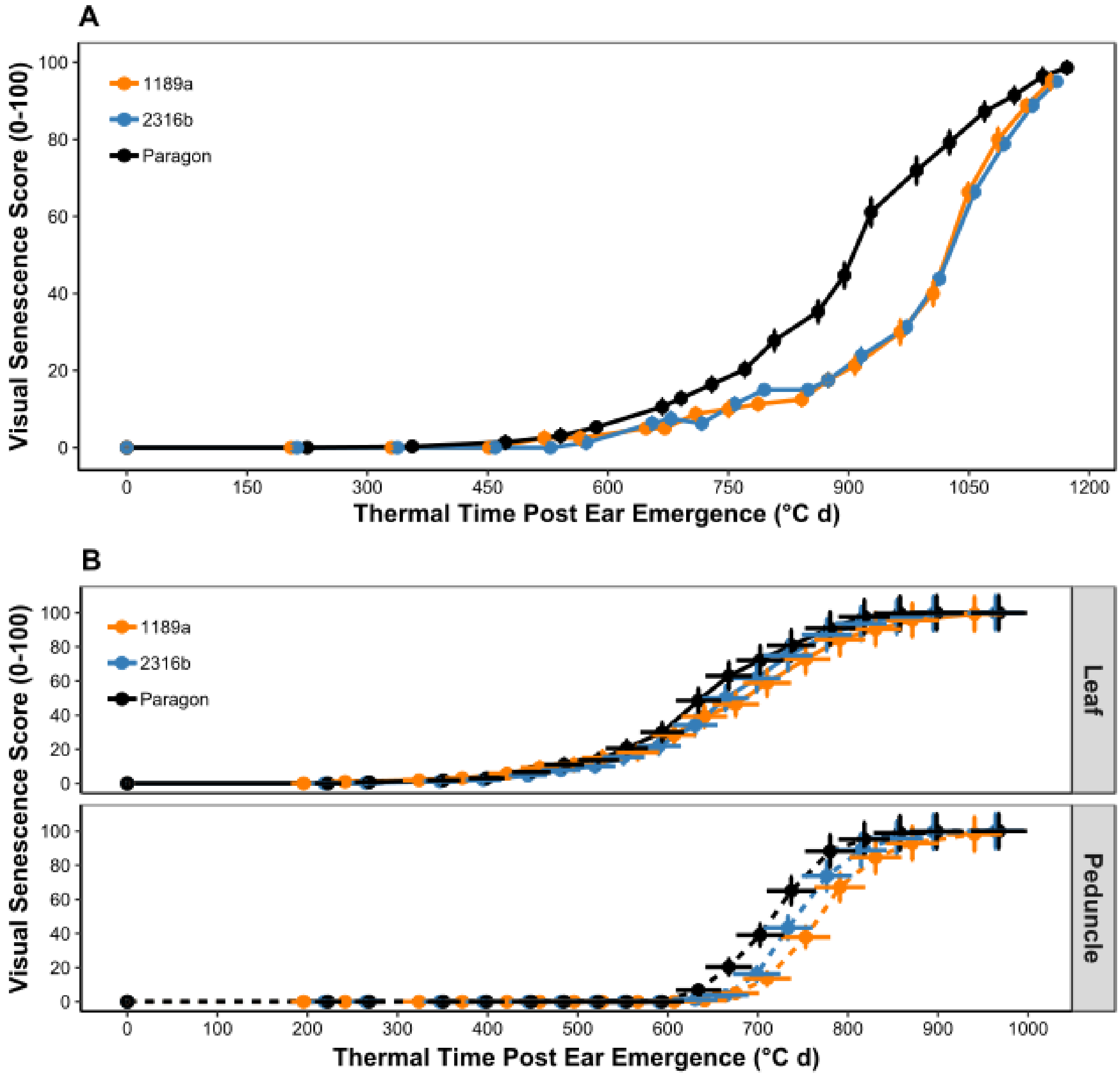

**Figure S2. Senescence phenotypes corresponding to lines used in grain filling experiments in 2018.** Progression of flag leaf **(A)** and peduncle **(B)** senescence for lines 1189a (orange, left) and 2316b (blue, right) is significantly delayed compared to cv. Paragon, P-value < 0.0001. Senescence and grain filling phenotypes mirror one another and relate to severity in onset. Senescence scored visually using a 0-100 scale 2-4 times per week from anthesis. Mean ± SD, n = 2.

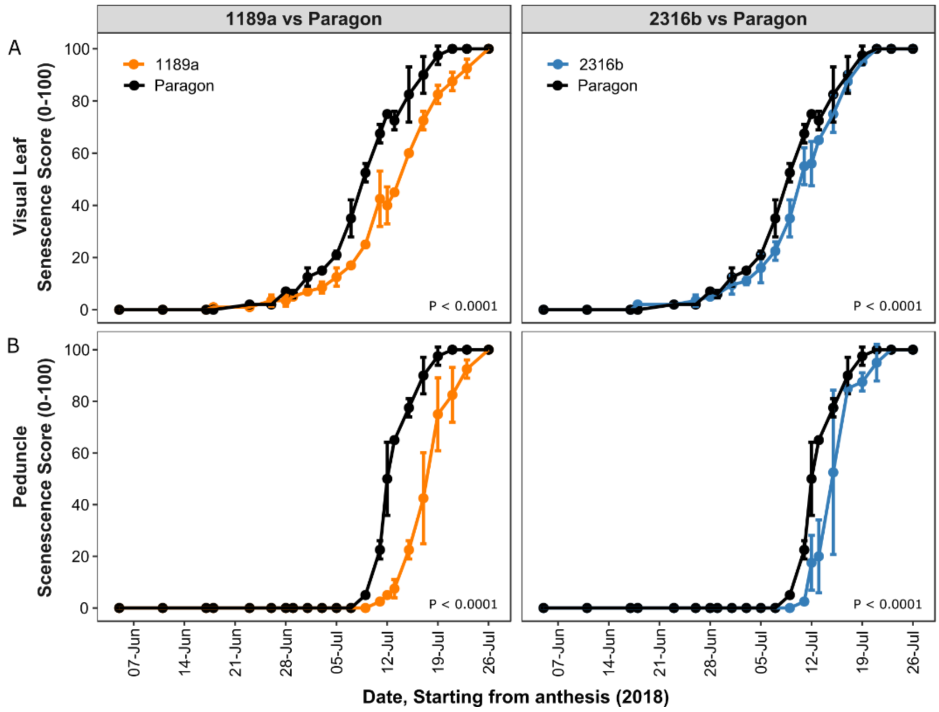

**Figure S3. Delayed senescence is associated with grain fill extension.** Results of grain filling experiments conducted for 1189a (orange, right) and 2316b (blue, left) in 2017. 37 daa grain moisture content of mutant lines remained elevated compared to cv. Paragon, P-values < 0.01 **(B)**. Frequency of which significant differences in grain moisture content are recorded reflect differences in senescence duration of 1189a and 2316b compared to cv. Paragon **(A)**. Differences in dry weight accumulation over time were significant between line 2316b and cv. Paragon, P-value < 0.01, but not 1189a, P-value > 0.1 **(C)**, and were unrealised regarding final grain weight. Senescence, grain moisture content (%) and dry grain weight (10 grains in mg) were recorded at 3 to 5-day intervals starting from anthesis. Mean ± SD, n = 4-5, 1 plot per line. Printed P-values represent overall differences (ANOVA). Pairwise differences indicated at corresponding time points, P-values: * < 0.05, ** < 0.01, *** < 0.001 (Tukey post-hoc test).

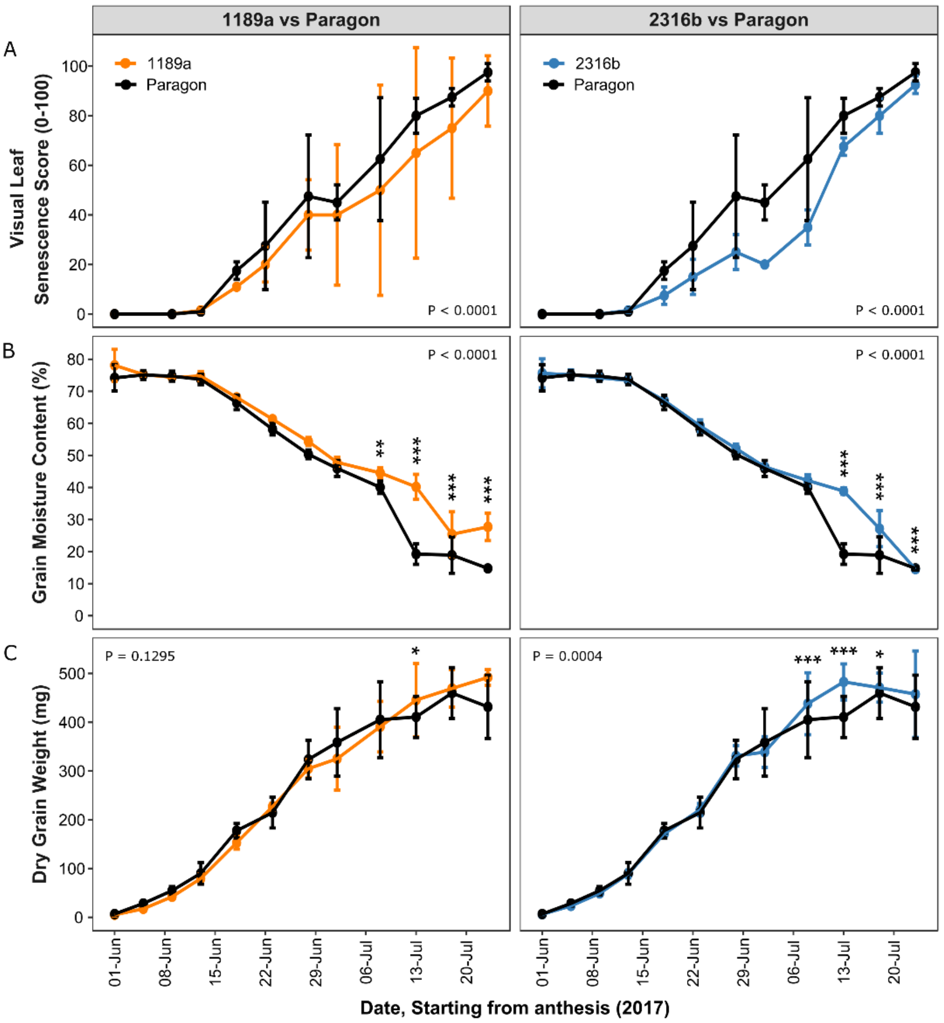

**Figure S4. TT70 scores for Paragon x 1189a $F_4$ RILs (2016 to 2018).** Senescence metric **'**Time to visual leaf senescence score 70 (°C d)' proved reliable in identifying senescence extremes for inclusion in bulks. Mean ± SD for each RIL in **(A)** 2016, n = 1; **(B)** 2017, n = 3; **(C)** 2018, n = 2. Coloured bars represent parents and RILs included in bulks, 1189a (orange), cv. Paragon (black), 'staygreen' (light orange, n = 17), 'non-staygreen' (light purple, n = 17).

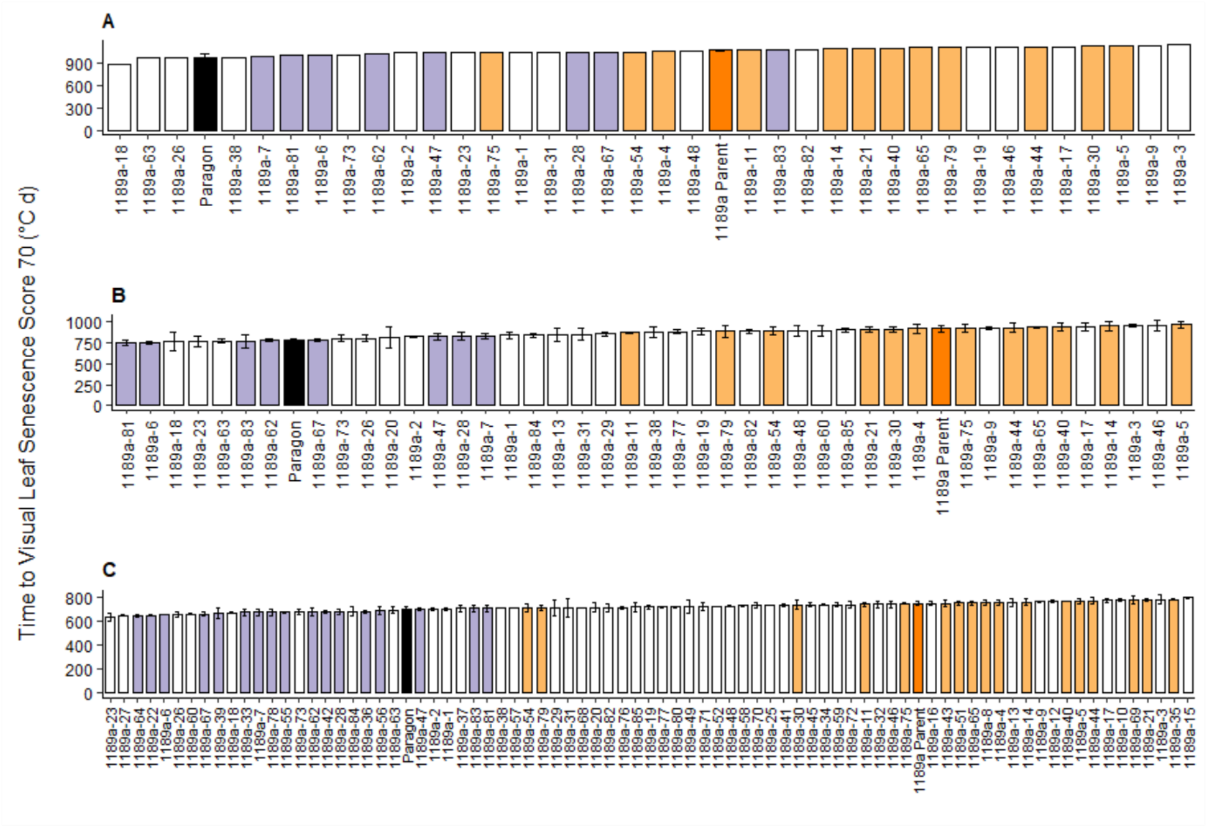

**Figure S5. TT70 scores for Paragon x 2316b $F_4$ RILs (2016 to 2018).** Senescence metric 'Time to visual leaf senescence score 70 (°C d)' proved reliable in identifying senescence extremes for inclusion in bulks. Mean ± SD for each RIL in **(A)** 2016, n = 1; **(B)** 2017, n = 3; **(C)** 2018, n = 2. Coloured bars represent parents and RILs included in bulks, 2316b (dark blue), cv. Paragon (black), 'staygreen' (light blue, n = 15), 'non-staygreen' (red, n = 12).

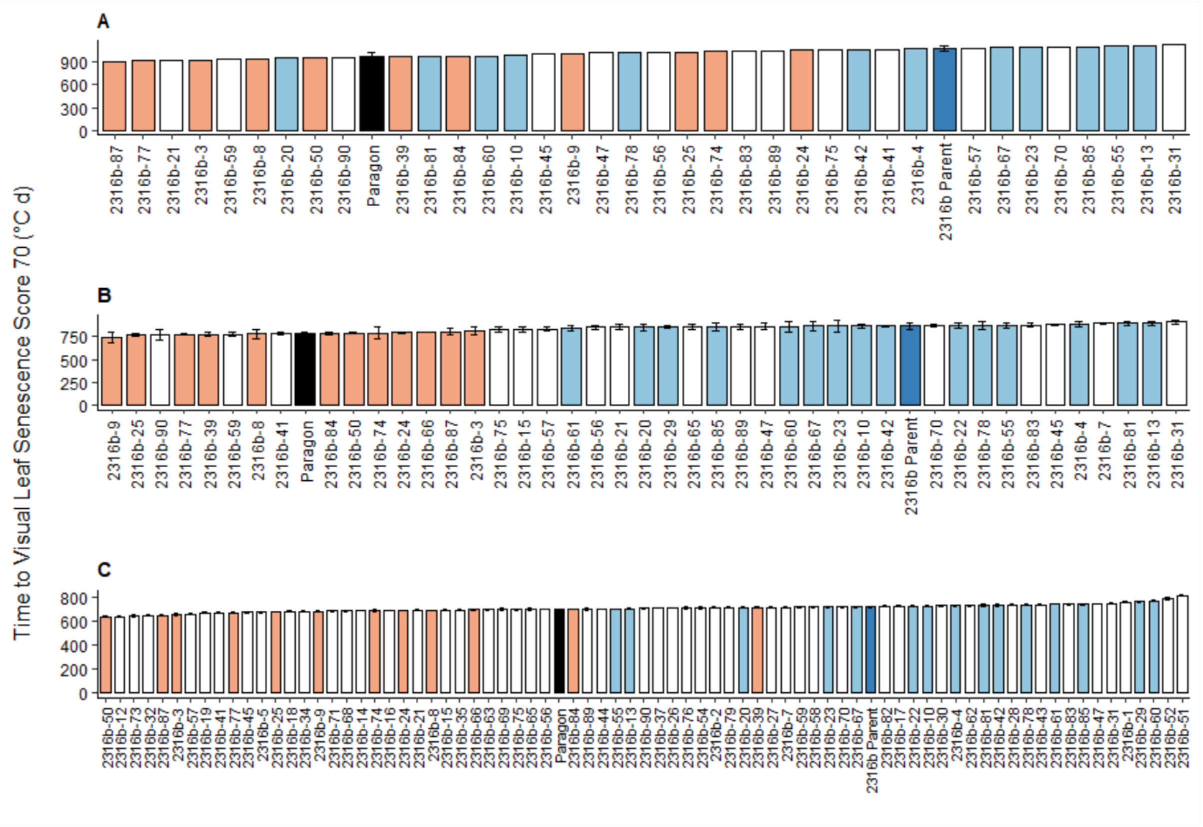


**Figure S6. Additional Phenotype x Genotype plots illustrating mode of inheritance.** Scatterplots displaying mean peduncle senescence values recorded in 2017 **(A)** and duration of leaf senescence (from GS55) recorded in 2018 **(B)** against *NAM-1* genotype. Paragon x 1189a $F_4$ RILs (orange, left), Paragon x 2316b $F_4$ RILs (blue, right). G/C cv. Paragon allele, A/T mutant allele. Phenotypic differences between contrasting homozygotes are significant, P-value < 0.001. Heterozygotes display intermediate to staygreen phenotypes. Black circle, genotypic mean ± SD. Differing numbers of RILs, including heterozygotes, grown each year, 2017, n = 43; 2018, n > 75.

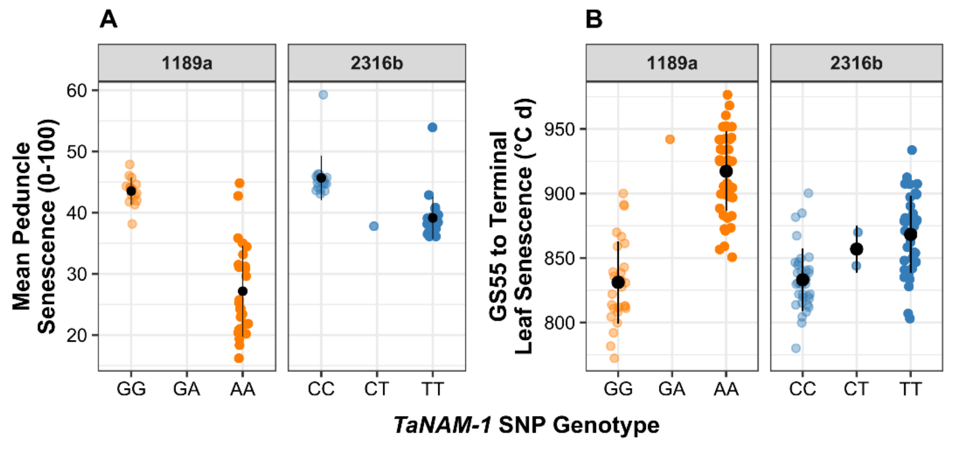

**Figure S7. Observation of co-dominant and additive modes inheritance for *NAM-1* mutations.** Senescence profiles of $F_3$ RILs heterozygous for *NAM-A1* and *NAM-D1* mutations (from independent crosses) support proposed differential modes of inheritance. Senescence profiles of homozygous RILs contrasting for *NAM-A1* were significantly different, P-value < 0.01, when grown in Cambridgeshire **(A)** and Norwich **(B)**, with heterozygotes resembling RILs homozygous for the *NAM-A1* mutation, P-value > 0.66. Phenotypic penetrance of the *NAM-D1* mutation is subject to greater environmental interaction. Differences in senescence progression of homozygous RILs contrasting for *NAM-D1* were significant when grown in Cambridgeshire, P-value < 0.05 **(C)**, not Norwich, P-value > 0.55 **(D)**. *NAM-D1* heterozygotes resemble RILs homozygous for the *NAM-D1* mutation, P-value > 0.9, and display intermediate to staygreen phenotype, supporting an additive mode of inheritance.

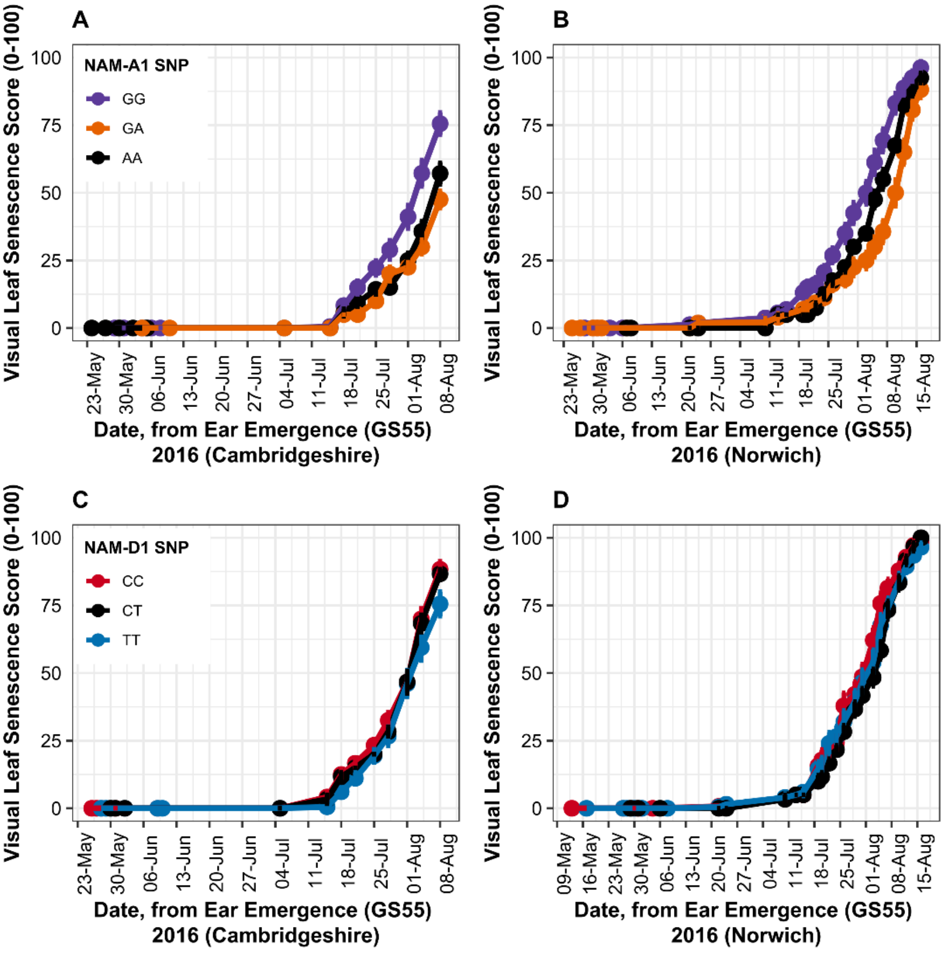

**Table S1.** *Differences in grain filling parameters correspond to severity in onset of senescence.*

Mean [$CI_{95\%}$], results of pairwise Tukey post-hoc tests following ANOVA. P-values, * < 0.05, ** < 0.01, *** < 0.001. Corresponding graphs, Supplemental Figure S3 (2017) and Figure 3 (2018).

| | | 2017 (dd/mm) | | | | 2018 (dd/mm) | | | |
|---|---|---|---|---|---|---|---|---|---|
| **Genotype** | **Grain Filling Component** | **08/07** | **13/07** | **18/07** | **23/07** | **12/07** | **17/07** | **20/07** | **23/07** |
| **Paragon** | **Moisture Content (%)** | **40.1 [37.7, 42.5]** | **19.2 [16.8, 21.6]** | **18.9 [16.5, 21.3]** | **14.8 [12.4, 17.2]** | **39.0 [37.5, 40.6]** | **16.2 [14.7, 17.8]** | **11.8 [10.2, 13.3]** | **8.2 [6.6, 9.7]** |
| | **Weight (10 grains in mg)** | **391 [370, 412]** | **420 [399, 441]** | **459 [438, 480]** | **438 [417, 459]** | **482 [436, 501]** | **423 [404, 442]** | **437 [418, 456]** | **443 [424, 462]** |
| **1189a** | **Moisture Content (%)** | **44.5 [42.1, 46.9] **** | **40.2 [37.8, 42.6] ***** | **25.4 [23.0, 27.8] ***** | **27.7 [25.3, 30.1] ***** | **42.6 [41.1, 44.2] **** | **35.3 [33.8, 36.9] ***** | **20.0 [18.5, 21.6] ***** | **14.4 [12.9, 16.0] ***** |
| | **Weight (10 grains in mg)** | **417 [396, 438]** | **452 [431, 473] *** | **480 [459, 501]** | **488 [467, 509]** | **473 [454, 492]** | **514 [494, 533] ***** | **476 [457, 496] **** | **506 [487, 525] ***** |
| **2316b** | **Moisture Content (%)** | **42.2 [39.8, 44.6]** | **38.9 [36.2, 41.6] ***** | **27.1 [24.7, 29.5] ***** | **14.5 [12.1, 16.9] ***** | **42.0 [40.5, 43.6] **** | **30.4 [28.8, 31.9] ***** | **16.4 [14.8, 17.9] ***** | **9.6 [8.0, 11.1]** |
| | **Weight (10 grains in mg)** | **453 [432, 474] ***** | **482 [460, 504] ***** | **497 [476, 518] *** | **462 [441, 483]** | **464 [444, 483]** | **466 [446, 485] ***** | **431 [412, 450]** | **423 [404, 442]** |

### Table S2. Exome Capture Coverage and identified SNPs

| Sample | No. of Positions | Average Coverage (%) | Positions Interrogated (QUAL ≥ 20) | Positions with ambiguous alleles | No. of SNPs | Transitions (G:A, C:T) | % |
|---|---|---|---|---|---|---|---|
| **1189a Staygreen bulk** | 131890905 | 41.87 | 87436528 | 281007 | 63667 | 20368 | 32.0 |
| **1189a Non-SG bulk** | 131808060 | 39.72 | 86231450 | 289595 | 62153 | 19840 | 31.9 |
| **2316b Staygreen bulk** | 133560822 | 43.16 | 89672224 | 277857 | 66506 | 21240 | 31.9 |
| **2316b Non-SG bulk** | 131470978 | 35.80 | 80600195 | 257888 | 57320 | 18437 | 32.2 |
| **1189a Parent** | 131990025 | 42.66 | 88654268 | 285206 | 65787 | 21503 | 32.7 |
| **2316b Parent** | 134157857 | 43.12 | 89314438 | 278525 | 68215 | 22125 | 32.4 |
| **Paragon** | 133301196 | 39.05 | 85580450 | 269711 | 62916 | 20149 | 32.0 |

#### Table S3. KASP primers for 1189a

| Chromosome | Position | Primer Name | Paragon (Hex) | 1189a Mutant (Fam) | Common | Cycling Conditions |
|---|---|---|---|---|---|---|
| chr6A | 25687659 | #61 | CCAGGGTAGTTATCCTTCTACc | CCAGGGTAGTTATCCTTCTACt | TTGTTGTGGAGCTGCGATA | 30-40 cycles 57/72°C |
| chr6A | 46782541 | 6A#47 | GCAATGTGGTGTCTGCGACg | GCAATGTGGTGTCTGCGACa | TACGAGCTGCGCAAGCAGT | 30-40 cycles 60°C |
| chr6A | 52406777 | 6A#55 | CAAATGACTGATGGGTACCTTc | CAAATGACTGATGGGTACCTTt | GCTAATGTTCCTTCCATAAGTG | 30-40 cycles 60°C |
| chr6A | 55741222 | bsa#6A#1 | GTTGCAACAAAATTAGTCCACAAg | GTTGCAACAAAATTAGTCCACAAa | GTGATGCCAATTCTCAAAACG | 30-40 cycles 57/72°C |
| chr6A | 57668862 | 6A#38 | CCTCGTCTATCTGGAATTTGg | CCTCGTCTATCTGGAATTTGa | ACGATGGTTTGAGTGGAGAG | 30-40 cycles 57°C |
| chr6A | 66431084 | 6A#35 | AAAGGCATAAATCTCTCCATGg | AAAGGCATAAATCTCTCCATGa | CGCAATGACAGGTTTGTGCAA | 30-40 cycles 60/72°C |
| chr6A | 67373981 | bsa#6A#2 | GCCTACGCCTTCGCCTTCg | GCCTACGCCTTCGCCTTCa | CATGGAGACGGCGAAGACC | 30-40 cycles 57/72°C |
| chr6A | 76523438 | 6A#7 | GGTCGACCACCGGCTTCAc | GGTCGACCACCGGCTTCAt | CTCAACCCTTACCTCGAGGC | 30-40 cycles 57°C |
| chr6A | 77099433 | bs#6A#3 | CTCGTGCATGATCCAGTTGg | CTCGTGCATGATCCAGTTGa | GTCAAGAAGGCGCTCGTCT | 30-40 cycles 57°C |
| chr6A | 80699949 | 6A#19 | TACAAGTGTATGCCTTTGCTGg | TACAAGTGTATGCCTTTGCTGa | GTAGTAAGTGCTATTATATAACCTA | 30-40 cycles 60°C |
| chr6A | 84072498 | bsa#6A#4 | GTAATCTTGCACCTATGTATAAg | GTAATCTTGCACCTATGTATAAa | GATCTAGCTAGCTTCTGCTG | 30-40 cycles 57°C |
| chr6A | 93079117 | 6A#24 | ATGAACAACAGTCGCCCTGc | ATGAACAACAGTCGCCCTGt | ACTGTCGAAAGTTATTGACTGT | 30-40 cycles 57°C |
| chr6A | 94068104 | bsa#6A#5 | AACACAAGAGATGGAGTGAGg | AACACAAGAGATGGAGTGAGa | AAGAAGGGGCAGTATTTTGATA | 30-40 cycles 57°C |
| chr6A | 96601065 | bsa#6A#6 | CGCCATGACGGCGATCGAg | CGCCATGACGGCGATCGAa | CCGAAGTGTGGACGAGGATC | 30-40 cycles 57/72°C |
| chr6A | 104309957 | 6A#30 | AAATGCAAGTGCTAGGTTGGg | AAATGCAAGTGCTAGGTTGGa | CTTCCTCTACACTCGGTTTCT | 30-40 cycles 57°C |
| chr6A | 105559955 | 6A#1 | GGTCGACCACCGGCTTCAc | GGTCGACCACCGGCTTCAt | CTCAACCCTTACCTCGAGGC | 30-40 cycles 57°C |
| chr6A | 116709274 | bsa#6A#8 | CTGAAGTAGCATGACAGCTGc | CTGAAGTAGCATGACAGCTGt | ACAATGCAAGTTCTATTGGTGT | 30-40 cycles 57°C |
| chr6A | 142783784 | 6A#12 | GAGCCTGGAAGTATTGCTGc | GAGCCTGGAAGTATTGCTGt | CCTTGGACAAGCCAACAATC | 30-40 cycles 57/72°C |
| chr6A | 379658960 | 6A#8 | AACAAAAGTCAACAGGTAACTAc | AACAAAAGTCAACAGGTAACTAt | GATGTCCCAAAGTTTACAAATC | 30-40 cycles 57°C |
| chr6A | 431054756 | 6A#4 | TCAGCTCGCCACATAGTCAg | TCAGCTCGCCACATAGTCAa | GCACGATCACTCAACATCCT | 30-40 cycles 57°C |
| chr6A | 459076102 | 6A#5 | CATCGCCCCGCGAAGCGg | CATCGCCCCGCGAAGCGa | GTCACCCGAAACCACCTGG | 30-40 cycles 57/72°C |
| chr6A | 479267518 | 6A#14 | ACCTTCCTCAACCTGCCAAc | ACCTTCCTCAACCTGCCAAt | GAAACAGGCATGATCAGGAG | 30-40 cycles 57/72°C |
| chr6A | 545816649 | 6A#GS2 | GCTTATGTCTTTGCGATCTGAg | GCTTATGTCTTTGCGATCTGAa | GACACACCAATGTGAGGAAAG | 30-40 cycles 57/72°C |
| chr6A | 549531562 | 6A#GS3 | GGTGGGTGTAGTTCATCTCTc | GGTGGGTGTAGTTCATCTCTt | TCACAAACGAGCTTTTCTCGG | 30-40 cycles 57°C |
| chr6A | 564876675 | #64 | ACATAGGAAGTGATATCTGTGg | ACATAGGAAGTGATATCTGTGa | AAGCAATCGTTCAAGTATGGC | Hydrocycler |
| chr6A | 581749266 | 6A#GS5 | AAAGGCAATGCAGCCACCAc | AAAGGCAATGCAGCCACCAt | CCTAAAGAGATTGCATGGAAGT | 30-40 cycles 57°C |
| chr6A | 585431913 | 6A#GS6 | CAGATACTGCATACTAGGATGg | CAGATACTGCATACTAGGATGa | AGGCGATTAGTAGTTTCACTC | 30-40 cycles 57°C |
| chr6A | 591732057 | 6A#GS8 | TGGCCAGCATGATGGCGATg | TGGCCAGCATGATGGCGATa | CTCTACCTCGGCTACTCAC | 30-40 cycles 57°C |
| chr6A | 603205645 | #63 | GTAAAGGACGGATTTATATACAGg | GTAAAGGACGGATTTATATACAGa | GTGATCATTCAAATGTGTCGGTG | Hydrocycler |

### Table S4. KASP primers for 2316b

| Chromosome | Position | Primer Name | Paragon (HEX) | 2316b Mutant (FAM) | Common | Cycling Conditions |
|---|---|---|---|---|---|---|
| chr6D | 36628312 | 2316b-6D#1 | CGAATTGACGCGATTTTTTCTc | CGAATTGACGCGATTTTTTCTt | GAGAGGAAAAGACGTCCGC | 30-40 cycles 57°C |
| chr6D | 42295410 | 2316b-6D#13 | CAATAATTAATCCGGGTCCATg | CAATAATTAATCCGGGTCCATa | CAAAATCTGACCAGTTGATATG | 30-40 cycles 57°C |
| chr6D | 50958137 | 2316b-6D#2 | GGCTTCAGCAGCAAATTCATAc | GGCTTCAGCAGCAAATTCATAt | CACTACTCCAGCAACTTCTTAG | 30-40 cycles 60°C |
| chr6D | 57044844 | 2316b-6D#14 | GGTCACCCGGGCGAAATGg | GGTCACCCGGGCGAAATGa | CCCCCATCATCCGGGATGA | 30-40 cycles 60°C |
| chr6D | 60487321 | 2316b-6D#15 | CGCTCGTGTTCTACCGCGg | CGCTCGTGTTCTACCGCGa | TCGGTGAGGCGGTATTCATG | 30-40 cycles 60°C |
| chr6D | 65341694 | 2316b-6D#16 | GAAGGTGCTTAAGACGCTGg | GAAGGTGCTTAAGACGCTGa | CACTACCACATCATCACTGTC | 30-40 cycles 57°C |
| chr6D | 72022762 | 2316b-6D#17 | GGATTGGTCTGCAAGAAAATAAg | GGATTGGTCTGCAAGAAAATAAa | TCAAATTTTGGTATGGAGCATTTT | 30-40 cycles 60°C |
| chr6D | 79076045 | 2316b-6D#3 | GTGGAGGACACGATGGCAg | GTGGAGGACACGATGGCAa | GAGATAGAACGGGGTCCTG | 30-40 cycles 57°C |
| chr6D | 84222670 | 2316b-6D#4 | GGAACTCAAGTTCAGGTAAAGg | GGAACTCAAGTTCAGGTAAAGa | ATCGGTCAGCTGAAAGCCC | 30-40 cycles 60°C |
| chr6D | 86733052 | 2316b-6D#18 | CAATCTGCTTCCATGCTACGg | CAATCTGCTTCCATGCTACGa | AATGCAACTTTTACAGCACAAAAT | 30-40 cycles 60°C |
| chr6D | 98263118 | 2316b-6D#5 | CACAGAATCTCCTCGGCGc | CACAGAATCTCCTCGGCGt | CAGAAACTAATTTGGACCACG | 30-40 cycles 57°C |
| chr6D | 110872402 | 2316b-6D#6 | GCAACAGCTCTCCCATCGc | GCAACAGCTCTCCCATCGt | CCAAGGATGCAGCCTCAGC | 30-40 cycles 57°C |
| chr6D | 119464714 | 2316b-6D#7 | GGTCCATCATTACAGGCCGc | GGTCCATCATTACAGGCCGt | TCGGCGCTCACAGCTGCAA | 30-40 cycles 60°C |
| chr6D | 139310981 | 2316b-6D#8 | CCACTAGACAGGTATATACCc | CCACTAGACAGGTATATACCt | CTGCAACAGCCGTTCAAAAG | 30-40 cycles 57°C |
| chr6D | 140102996 | 2316b-6D#9 | GTGTTATTCAGGGCGAAACGg | GTGTTATTCAGGGCGAAACGa | GGTCAACTGCTCTATGCTGC | 30-40 cycles 60°C |
| chr6D | 148079666 | 2316b-6D#10 | GGACACCGACATGCCGGg | GGACACCGACATGCCGGa | TGCACGAGCGGATAGCCAA | 30-40 cycles 57°C |
| chr6D | 264681596 | 2316b-6D#11 | TTTTTAGCAACTGCTGCATTGc | TTTTTAGCAACTGCTGCATTGt | GCTATTACCATTCACATCCTG | 30-40 cycles 57°C |
| chr6D | 292379919 | 2316b-6D#12 | TGAAACCTGACGGAAGAAAGg | TGAAACCTGACGGAAGAAAGa | TCTATATACTGGCAAAGCAGC | 30-40 cycles 57°C |
| chr6D | 352357451 | 2316b-6D#19 | GGTGGTCGGCGAGGCGAg | GGTGGTCGGCGAGGCGAa | CTTCGCCATCCTCCACCAC | 30-40 cycles 57°C |
| chr6D | 363004043 | 2316b-6D#20 | CGCTGCGGTCCGAGGGg | CGCTGCGGTCCGAGGGa | CTGGGCCCCGCACGTGA | 30-40 cycles 57°C |
